# Cell fate plasticity of xylem-pole-pericycle in *Arabidopsis* roots

**DOI:** 10.1101/2024.06.17.599297

**Authors:** Xin Wang, Lingling Ye, Jing Zhang, Charles W. Melnyk, Ari Pekka Mähönen

## Abstract

In *Arabidopsis* roots, xylem-pole-pericycle (XPP) cells exhibit remarkable cell fate plasticity by contributing to both lateral root (LR) and cambium formation. Despite significant progress in understanding these individual processes, the mechanism orchestrating these two fates and their effects on root architecture and secondary growth remain unclear. Here we combined lineage tracing with molecular genetics to study the fate dynamics of XPP cells. We showed that developmentally arrested lateral root primordia (LRP) that fail to emerge as lateral roots, gradually obtain cambium identity thus contributing to secondary growth. Conversely, preestablished procambium identity within XPP cells can be reverted to LR identity when simulated by auxin, an important player in LR development. This competence for auxin-induced LR formation from XPP cells, termed LR potency, however, decreases as the root matures. We found key cambium regulators play essential roles in shaping LR potency by promoting cambium activation and inhibiting LR development. Consistently, corresponding mutants with impaired cambium activity display broader LR potency. Moreover, cytokinins, essential players in cambium development, facilitate the identity transition of LRPs to cambium and reduce LR potency through key cambium regulators. Overall, our findings highlight the inherent cellular plasticity of XPP cells and elucidate how plant hormones influence root architecture and secondary growth through balancing the two cell fates of XPP cells.

## Introduction

Plants, as sessile organisms, have developed adaptive abilities to thrive in challenging environments. An exemplary demonstration of plant developmental plasticity lies in the versatility of xylem-pole-pericycle (XPP) cells and their various destinies. Within the root tip of *Arabidopsis*, a cluster of XPP cells is designated as prebranch sites, primed for subsequent formation of lateral root primordium (LRPs) ^1-4^. During secondary growth, XPP cells not participating in LR formation adopt the identity of cambium (referred to as cambium-fate XPP cells), contributing to secondary growth^5^. Lineage tracing studies have shown that XPP cells give rise to both vascular cambium, a key driver of secondary growth, and cork cambium, which produces the protective periderm layer^5^. Additionally, overlapping molecular mechanisms regulating pluripotent callus induction and LR development further underscores the high plasticity of XPP cell fate^6-8^.

The key plant hormones, auxin and cytokinin, play antagonistic roles in LR development. Auxin signaling influences every stage of LR development, including prebranch site formation, founder cell specification, LR initiation, as well as the subsequent patterned division and differentiation of LR primordia (LRP), culminating in the emergence of LRs by penetrating through overlying tissues^9-11^. Conversely, cytokinins impede LR initiation, proper positioning along the primary root, and the development^10,12-17^. While cytokinins exhibit inhibitory effects on LR development, they promote the activation of the cambium and serve as rate-limiting regulators of cambial activity and secondary growth^18,19^. Cytokinin treatment can prematurely activate cambium for secondary growth^19^. Therefore, while auxin promotes XPP cells to develop into LRs, cytokinins promote cambium formation. However, the mechanisms to balance between these two destinies are poorly understood.

LR initiation occurs periodically in the root tip region. Intriguingly, LRPs at various developmental stages are not arranged chronologically along the root. Instead, younger LRPs often present between two developmentally more advanced LRs/LRPs^20,21^. A prior investigation employing an enhancer trap line revealed that a considerable proportion of developmentally arrested LRPs fail to emerge later^20^, leaving their ultimate fate unknown. While it is known that auxin can induce *de novo* LR formation from XPP cells in a manner independent of the prebranching mechanism^2,22^, the competence for auxin-induced LR formation, termed LR potency here, and its potential conflict with cambium identity acquisition during secondary growth, remain to be investigated.

In this study, assisted by novel lineage tracing systems, we demonstrated that a significant portion of slowly developing LRPs fail to progress into emergence and instead, enter a silent state. These developmentally arrested LRPs eventually relinquish their original LR identity and adopt cambium identity. Conversely, the pre-established procambium identity within XPP cells can be reverted into LR identity stimulated by auxin. As the root matures, the LR potency decreases as a result of inhibitory effect of key cambium regulators. Cytokinins, consistent with their key roles in cambium development, promote the arrest of young LRPs and facilitate the transition from LR identity to cambium identity. Cytokinins also lower auxin-mediated LR potency via key cambium regulators. These findings not only underscore the remarkable cellular plasticity of XPP cells, but also revealed how auxin and cytokinins balance these two cell fates of XPP cells, thereby influencing root architecture and secondary growth.

### Developing CRE-lox based LR-tracing systems

Arrested LRPs are challenging to detect and track, especially in the more mature part of root (Fig. 1a). To overcome this, we developed an inducible CRE-lox-based LR-tracing system. We first assessed two LR-specific promoters, *pGATA23*^23^ and *pHB53*^24^, and made estradiol inducible versions of them, *pGATA23-XVE* and *pHB53-XVE*. We fused each inducible promoter with the CRE recombinase coding sequence and introduced the constructs into a prescreened *35S-Loxp-erGUSpYFP* background (Fig. 1b). The erGUSpYFP fusion contain ER-localized GUSplus reporter fused with YFP reporter thus allowing flexible analysis of the clones. In T1 transgenic plants, we observed that after a 1-day estradiol treatment, YFP-labeled LR/LRP clones were efficiently induced on the convex side of the root curve. Comparative analysis revealed that, in contrast to *pHB53*, the GATA23 inducible promoter induced continuous YFP signals in the lower part of the root, resembling its native promoter’s expressing pattern^23,25^. Consequently, *pHB53-XVE* exhibited more specific labeling of LRs/LRPs compared to *pGATA23-XVE* (Fig. 1c).

**Fig. 1:**
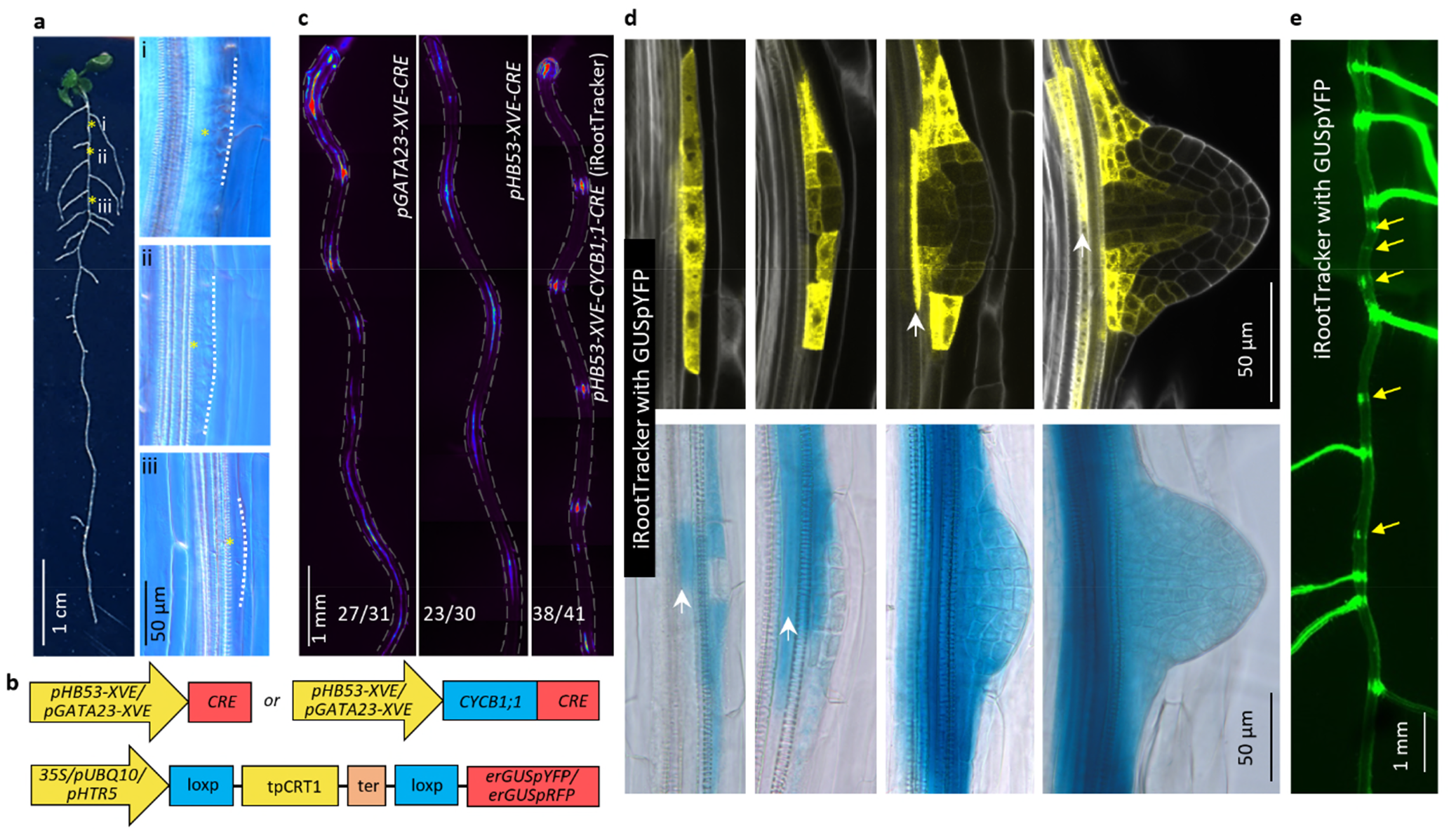
Establishment of CRE-lox based LR-tracing systems. **a**, left panel, an overview of root architecture of an 8-day-old seedling with arrested LRPs. Right panel, a close-up view of developmentally arrested LRPs, outlined with white dashed lines. Yellow asterisks indicate relative positions of arrested LRPs in the root, **b**, A schematic of LR-tracing systems. CRE recombinase variants are induced with two different inducible, LR-specific promoters. CRE catalyzes recombination between two loxp sites leading to constitutive expression of reporters driven by *35S, pUBQlO* or *pHTR5*. **c**, A comparison of distinct LR-specific inducible promoters and CRE recombinase variants. Three-days old T1 seedlings were transferred to induction medium for 1 day before stereo microscopy. Numbers represent the frequency of the observed types of YFP clones in independent T1 samples. Roots are outlined with grey dashed lines. Combination of *pHB53-XVE-CYCBl;l-CRE* and *35S-Loxp-erGUSpYFP*, named iRootTracker, was used in the subsequent experiments, **d**, In the iRootTracker, a detailed confocal inspection and GUS staining revealed that LR/LRPs are faithfully marked during their morphogenesis. White arrows indicate unspecific clones associated with overlying LRs/LRPs. **e**, An example showing arrested LRPs (yellow arrows) in the upper region of a 10-days old root, as visualized by the iRootTracker under a fluorescent stereo microscope. Three-days old seedlings were first given 1 day induction then transferring to estradiol-free medium for another 6 days.

Despite being more specific for LRs/LRPs, YFP clones generated by *pHB53-XVE* marked a relatively broad domain beyond the LR/LRP itself (Fig. 1c). We then tested the CYCB1;1-CRE, to restrict the recombinase activity into G2/M phase of cell cycle (Fig 1b) ^5,26^. Compared with CRE, the CYCB1;1-CRE enabled the generation of smaller yet LR/LRP-specific clones (Fig. 1c). Therefore, the CYCB1;1-CRE recombinase was used for all further studies. Detailed confocal examination revealed nearly 100% (359/360 clones in 38 roots) of YFP clones specifically marked LRs/LRPs, with over 80% (250/298 clones in 36 roots) detected LRs/LRPs exhibiting YFP signal (Fig. 1d). GUS reporter assay showed a similar result (Fig. 1d). With such an inducible LR-tracing system, the arrested LRPs can be easily detected and tracked just under a fluorescent stereo microscope (Fig. 1e). We named this inducible LR-tracing system as iRootTracker.

In addition, by utilizing the HB53 native promoter, we also established a stable LR-tracing system (named RootTracker) and validated its reliability in LR/LRP tracing (Supplementary Fig. 1). To demonstrate versatility of this system, we grew the RootTracker line under different nutritional, hormonal or stress conditions (Supplementary Fig. 2). In addition to expected root architecture phenotypes^27-29^, the LRs/LRPs under each condition were clearly labeled with YFP, indicating the wide compatibility of this system with various growth conditions (Supplementary Fig. 2). With the RootTracker, the root architecture can be easily visualized and quantified under a fluorescent stereo microscope, significantly simplifying LR-related quantification compared to conventional methods. For example, quantifying the LR/LRP number within a fixed region of a 7-day-old root took about 25s with the RootTracker, while for the same region of the same root, standard microscopy required more than 5 minutes, without including time spent for sample preparation (Supplementary Fig. 3a). Implementation of such a system also improves the accuracy of LR/LRP quantification: an average of 30 LR/LPRs could be detected using the RootTracker under fluorescent stereo microscope, compared with 25 LR/LRPs with standard microscopy (Supplementary Fig. 3b).

Occasionally, we observed unspecific induction of YFP/GUS in the vascular cells underneath a LR/LRP in both systems (Fig. 1d and Supplementary Fig. 1). However, this did not affect LR/LRP identification or quantification as these non-specific expressions were consistently associated with the overlaying YFP/GUS-marked LRs/LRPs. We also noticed that LRs/LRPs were not always fully marked by YFP/GUS expression (Fig. 1d and Extend Data Fig. 1). This might be due to either loss of recombination events in some cells of a LRP, or by the inherent property of the 35S promoter, which drives the YFP/GUS expression after the recombination. Additionally, the youngest LRPs near the root tip exhibited weak signal, likely because these LRPs might not have enough time to accumulate sufficient amount of YFP/GUS for visualization after recent recombination events (Supplementary Fig. 1). Thus, we typically quantified LR/LRPs within a fixed region of the root rather than the entire root. In T2 generation of the RootTracker lines, we also noticed a small portion of seedlings (ranging from 0.3%-2.8% for different lines) showing YFP expression in the whole seedling, indicating unspecific recombination events in the germ cell line. In conclusion, iRootTracker and RootTracker facilitate convenient and high throughput, real-time LR development observation and quantification in an intact root, just under a fluorescent stereo microscope.

### Arrested LRPs eventually acquire cambium identity

Taking advantage of the iRootTracker, we could monitor each LRP development over an extended period. We first gave a 1-day estradiol treatment to 3-day-old seedlings to induce YFP expression in LRPs, then transferred the seedlings to estradiol-free medium and tracked each YFP-marked LRP (in the upper root region of about 1 cm) over the following days, by using a fluorescent stereo microscope (Fig. 2a-c). Our findings revealed that LR emergence predominantly occurred between day 5 and day 7 in this root region (Fig. 2c). At day 7, approximately half of all LRPs failed to grow out (Fig. 2c). Intriguingly, the YFP sectors originating from these developmentally arrested LRPs gradually expanded radially along with the overall secondary growth (Fig. 2d). Further examination post-GUS staining revealed a morphological shift of these LRPs from cubic shape to long, thin cambial-like cells, aligning their growth with the general secondary growth pattern (Fig. 2e). Additionally, within the same observation region, the auxin responsive reporter DR5 exhibited low expression in most LRPs, suggesting reduced competence for continuing LR development (Fig. 2f).

**Fig. 2:**
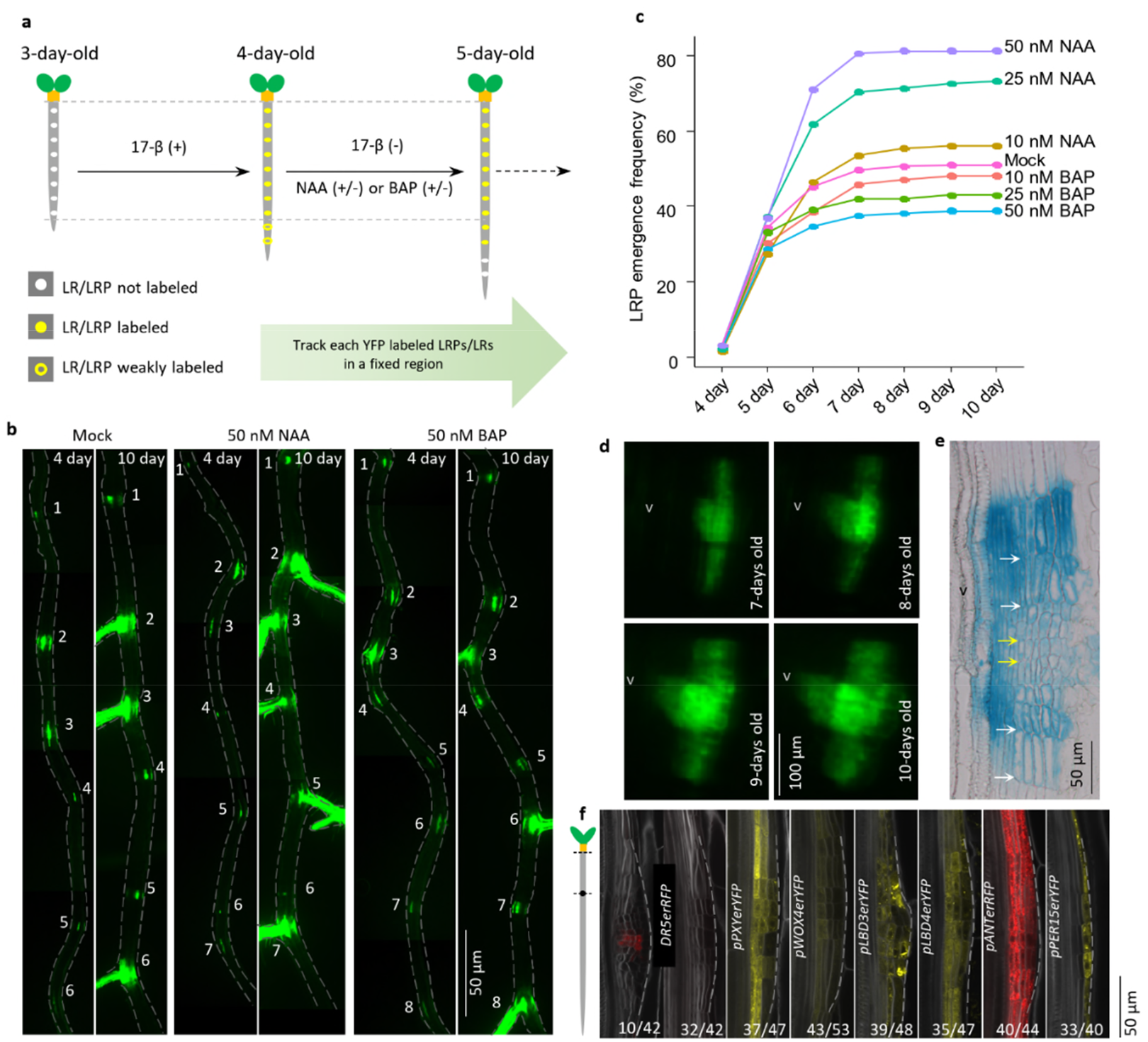
Arrested LRPs gradually acquire cambium identity. **a**, A schematic showing the strategy used for LR/LPR tracing with the iRootTracker. Three-day-old seedlings were first induced for 1-day to initiate LR-tracing before transferring them to estradiol-free medium with/without phytohormones. After transferring, each LR/LRP was tracked under a fluorescent stereo microscope until the roots were 10-days old. Note after a 1-day induction, the LRPs in the newly formed root region showing initially weak fluorescence (marked by yellow ring) were not followed. Therefore, a fixed root region marked between dashed lines was traced, **b**, Examples showing LR/LRP tracing in each root (outlined with grey dashed lines) under various conditions. In each panel, the same number labels the same LR/LRP when the root was 4-days old (left) and 10-days old (right). Roots are outlined with grey dashed lines, **c**, Quantifications of LRP emergence frequency over time in the presence of different concentrations of auxin (NAA) or cytokinin (BAP). Auxin promotes and cytokinins inhibit LRP outgrowth in a dose-dependent manner. The data are shown as mean values calculated from a total LRP number of 170 to 313, based on 22 to 44 individual roots. The raw data are available in the source data file, **d-e**, Lateral view of an arrested LRP showing radially expanded YFP sector over time (**d**) and GUS sector in a 2-weeks old root (e). Note the shorter cells (yellow arrows) in the region that used to be a LRP compared to the longer cells (white arrows) in regions below and above (**e**). V marks the vasculature region, **f**, While cambium markers showed their expression in arrested LRPs (outlined with grey dashed lines), DR5 expression was low, in the mature region of 7-day old roots. The left schematic shows the region of the root where imaging was carried out. When the roots were 3-days old, their root tips were labeled and 4 days later, only the root region above this label were sampled for confocal observation. Numbers refer to the frequency of observed phenotypes.

Lineage tracing and histological analysis suggest the arrested LRPs gradually acquire cambium identity to participate in the overall secondary growth. To validate this, we examined the expression of key vascular cambium regulator genes in LRPs, including the receptor kinase *PHLOEM INTERCALATED WITH XYLEM/TRACHEARY ELEMENT DIFFERENTIATION INHIBITORY FACTOR RECEPTOR* (*PXY/TDR*) ^30,31^, transcription factors *WUSCHEL RELATED HOMEOBOX 4* (*WOX4*) ^32^, *LATERAL ORGAN BOUNDARIES DOMAIN 3* (*LBD3*) and *LBD4*^19^, as well as *AINTEGUMENTA* (*ANT*) ^33^. Additionally, we also included a cork cambium reporter driven by *PEROXIDASE15* (*PER15*) promoter^34^. While expression of most of these reporters were absent in the developing LRPs (See below, Fig. 3), in the upper region of 7-day-old roots, most LRPs had obtained the expression of these vascular cambium reporters in the basal part of the primordia, adjacent to the protoxylem, as well as the PER15 expression in the outmost layer, indicating a transition from LR identity to cambial identity (Fig. 2f).

Next, we studied the mechanisms by which LRPs get arrested and obtain cambium identity. Given the importance of auxin gradient in LR development^15,35^, we proposed that the failure to maintain an adequate auxin gradient in some LRPs may result in arrest. Supporting this, our lineage tracing studies with the iRootTracker indicated that external auxin (naphthalene-acetic acid, NAA) supplementation to 4-day-old roots accelerated LRP outgrowth and reduced the number of arrested LRPs (Fig. 2a-c). Moreover, despite slowly acquiring cambial identity, a considerable portion (up to 20% depending on auxin levels) of arrested LRPs from 7-day old roots could still be reactivated by additional auxin supplementation, emphasizing the cell fate plasticity of XPP cells and their derivatives in response to internal and external stimuli (Supplementary Fig. 3c).

Arrested LRPs appeared to show increased resistance to auxin-triggered reactivation as the root matures. For example, relatively low levels of auxin (50 nM NAA) could efficiently reduce arrested LRP numbers when applied to a 4-day-old root (Fig. 2b, c), but only reactivated a few of arrested LRPs of a 7-day-old root. In the latter case, higher levels of auxin (up to 5 M) were required (Supplementary Fig. 3c).

### Cytokinins promote LR to cambium identity transition

Cytokinins are known to inhibit LR initiation and development^10,12-17^. Utilizing the iRootTracker, we demonstrated that cytokinin treatment induces arrest of existing LRPs in a dose-dependent manner (Fig. 2a-c). Since cytokinins also promote cambial activation and secondary growth^18,19^, we investigated whether cytokinin treatment facilitates the transition of LRPs to cambium. Following a 2-day BAP treatment in 3-day-old seedlings, we observed morphological changes of LRPs from a domed to flattened shape, accompanied with the emergence of expression of cambium reporters, including both vascular and cork cambium reporters, which were previously inactive in LRPs (Fig. 3). Unlike the other reporters, *LBD4* exhibited a heterogeneous expression across LRPs in the absence of cytokinin, with stronger expression in the boundary domain. However, following BAP treatment, this expression pattern shifted to a uniform expression in the basal domain of LRPs. Concurrently, LR and root meristem reporters tended to lose their expressions in LRPs (Fig. 3). Thus, cytokinin promotes LRP arrest and facilitates the acquisition of cambium identity.

**Fig. 3:**
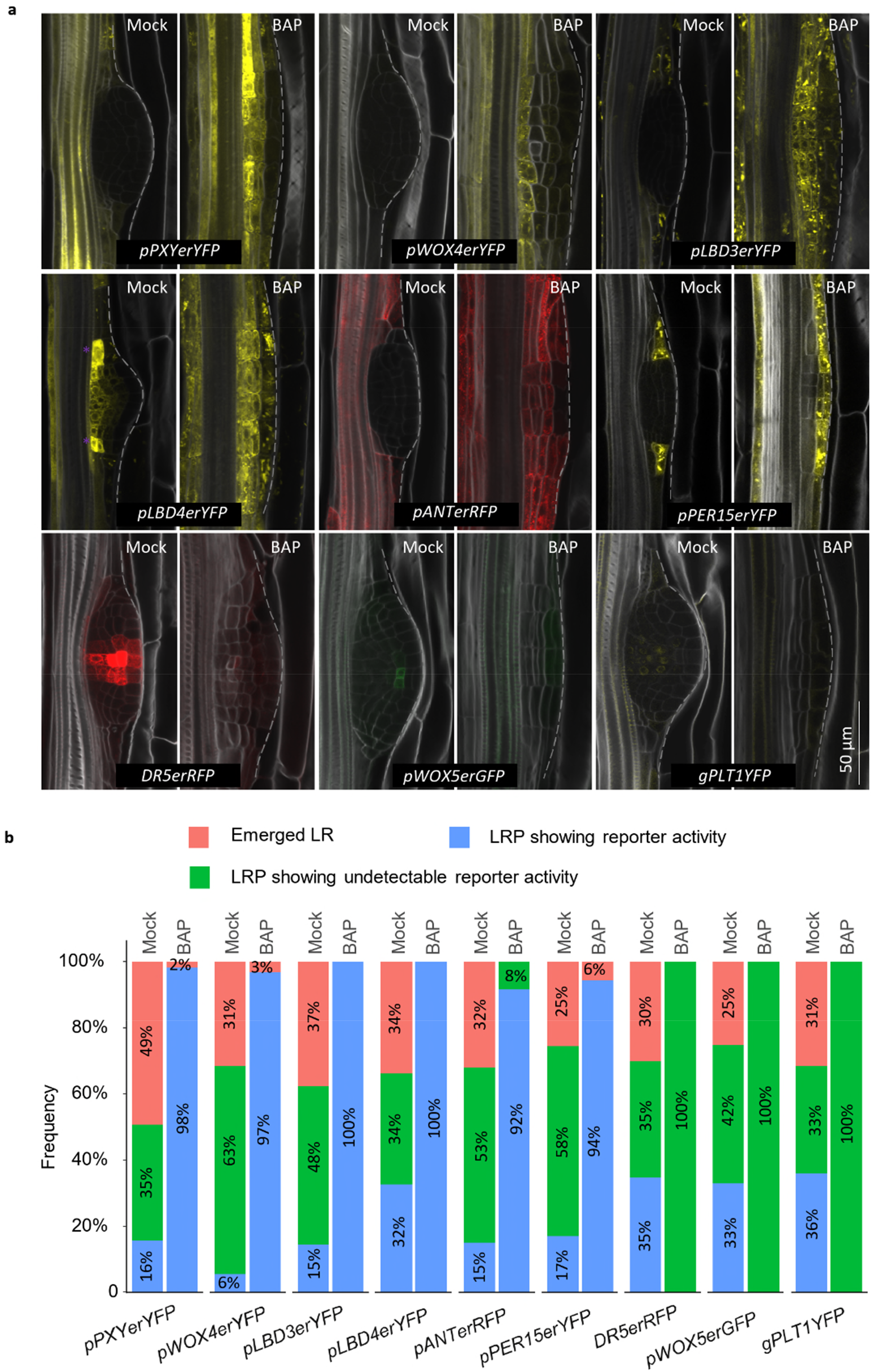
Cytokinins promote cambium identity acquisition of LRPs. **a**, Existing LRPs (outlined with grey dashed lines) acquired cambium identity with dampened expression of auxin and root meristem reporters, after a 2-day 1 µM BAP treatment in 3-days old seedlings. The LRPs from the newly grown part of root after a 2-day growth were not included. **b**, Statistics of distinct reporter expression in LRPs with/without BAP treatment. Note that *LBD4* was expressed in the subdomain of LRPs (classified into the category “LRP showing undetectable reporter activity”) in the mock treatment and throughout LPRs after BAP treatment **(a)**. The boundary expression of *LBD3* and *PER15* in some LRPs with the mock treatment **(a)** were classified into the category “LRP showing undetectable reporter activity”. The frequency is calculated from a total lateral organ number of 45 to 58, based on 12 to 16 individual roots. Raw data are available in the source data file.

In the 5-day-old roots with mock treatment, we found a small subset of LRPs (less than 17%) showing expression of cambium regulators (except *LBD4*) and twice as many LRPs with weakening DR5 reporter activity (about 35%), suggesting that arrested LRPs acquire cambium identity after auxin signaling has dampened (Fig. 3b and Supplementary Fig. 4a). Developmental arrest of LRP appears to take place during early stages, as suggested by their morphology (Supplementary Fig. 4a). LRPs in the 3-day-old roots seem to be highly sensitive to cytokinin treatment, as indicated by efficient cytokinin-mediated LR to cambium identity transition (Fig. 3). Similarly, a previous report has shown that younger LRPs exhibit greater sensitivity to cytokinin compared to developmentally more advanced LRPs^15^. It is possible that in younger LRPs, the auxin maximum has not been stabilized, making them more susceptible to external interference. We also found that a 10-h BAP treatment was insufficient to trigger *PXY/TDR, WOX4* and *ANT* expression in the most LRPs of 3-day-old seedlings, except for *LBD3*, which is a direct target of cytokinin signaling^19^ (Supplementary Fig. 4b). For *PXY/TDR*, for example, a 24-h treatment was required to induce its expression in LRPs (Supplementary Fig. 4c). In contrast, it has been shown cytokinins are able to rapidly (about 1.5 h) deplete the auxin efflux carrier PINFORMED 1 (PIN1) from the plasma membranes through modulating PIN1 endocytic recycling and thus redirect auxin flow and inhibit LR organogenesis^13,14^. It is thus likely that cytokinins promote the loss of LR identity and the acquisition of cambium identity in a sequential manner.

Altogether, our findings highlight the cell fate plasticity of LRPs. When LRPs enter arrested state, whether under normal conditions or due to cytokinin treatment, they undergo a transition to acquire cambium identity. This transformation enables them to contribute to root secondary growth, thereby serving as an additional source for cambium, alongside the cambium-fate XPP cells.

### Cambium regulators shape LR potency

The findings above demonstrated the reversibility from LRP identity to cambium identity. Previous reports indicate that XPP cells maintain the competence to form new LRs when stimulated by auxin, but the competence decreases in the segmented upper region of 10-day-old roots growing on medium containing 10 M of the auxin transport inhibitor 1-naphthylphthalamic acid (NPA) ^22,36^. To understand auxin-mediated LR potency in intact roots at a higher resolution, we characterized LR potency in 7-day-old roots by using the RootTracker with various supplemented auxin levels. We observed that 50 nM NAA triggered *de novo* LR formation in the region primarily near the root tip; while above this region, only the development of existing LRs/LRPs were affected (Supplementary Fig. 5a). When supplying 1 M NAA, after a 3-day induction, *de novo* LR formation was observed in a broader root region in the lower part of the root. In the upper part of the root, LR initiation seemed to occur as well, however, in most cases, it did not lead to LR outgrowth (Supplementary Fig. 5a). We then increased NAA level to 5 M and found massive and continuous *de novo* LR formation in the root region from about 1.5 cm below the hypocotyl towards the root tip, while the upper root region (about 1 cm) remained less responsive despite of such a high level of auxin (Supplementary Fig. 5a). These results indicated that LR potency is determined not only by the supplied auxin levels but also by the status of root maturity. Decreased auxin-induced LR potency as root matures was also observed by a detailed quantification of LR distribution in WT roots after a 3-day 5 M NAA treatment (Supplementary Fig. 5b). We also developed an XPP lineage tracing system by using an XPP-specific inducible promoter^37^ driving CYCB1;1-CRE expression. With this system we confirmed first that the newly formed LRs originated from XPP cells or their derivatives, and second, the LR potency decreased as root matures (Supplementary Fig. 5c). *ipt1;3;5;7*, which harbors loss-of-function mutations in four *ISOPENTENYL TRANSFERASE* (*IPT*) genes for cytokinin biosynthesis, lacks cambium activity in the root^18^. However, the mutant root exhibited exceptional LR potency, with auxin treatment triggering LR induction along the entire root length (Supplementary Figs. 5b and 6), thus confirming a previous finding^15^. The decreased LR potency within the WT roots and the differences in LR potency observed between WT and *ipt1;ipt3;ipt5;ipt7*mutant roots suggest the presence of a regulatory mechanism associated with secondary growth governing LR potency.

We proceeded to investigate the mechanism underlying the regulation of LR potency. Since cambium-fate XPP cells are destined for cambium formation, we proposed that the gradual acquisition of cambium identity may account for LR potency shaping. To test this, we first performed serial cross-sections to 7-day-old WT roots to explore the relationship between LR potency and status of maturity within XPP cells and their derivatives. Statistical analysis showed that cambium activation (that is, initiation of the periclinal cell divisions) predominantly occurred within the region spanning 1-1.5 cm below the hypocotyl of 7-day-old seedlings. Above this region, cambium was fully activated, while below it, cambium activation rarely occurred (Supplementary Fig. 6). This cambium activation pattern aligned well with LR potency along the root: extensive LR induction was observed below this region (Fig. 4a and Supplementary Figs. 5b and 6), while LR induction was limited above it.

**Fig. 4:**
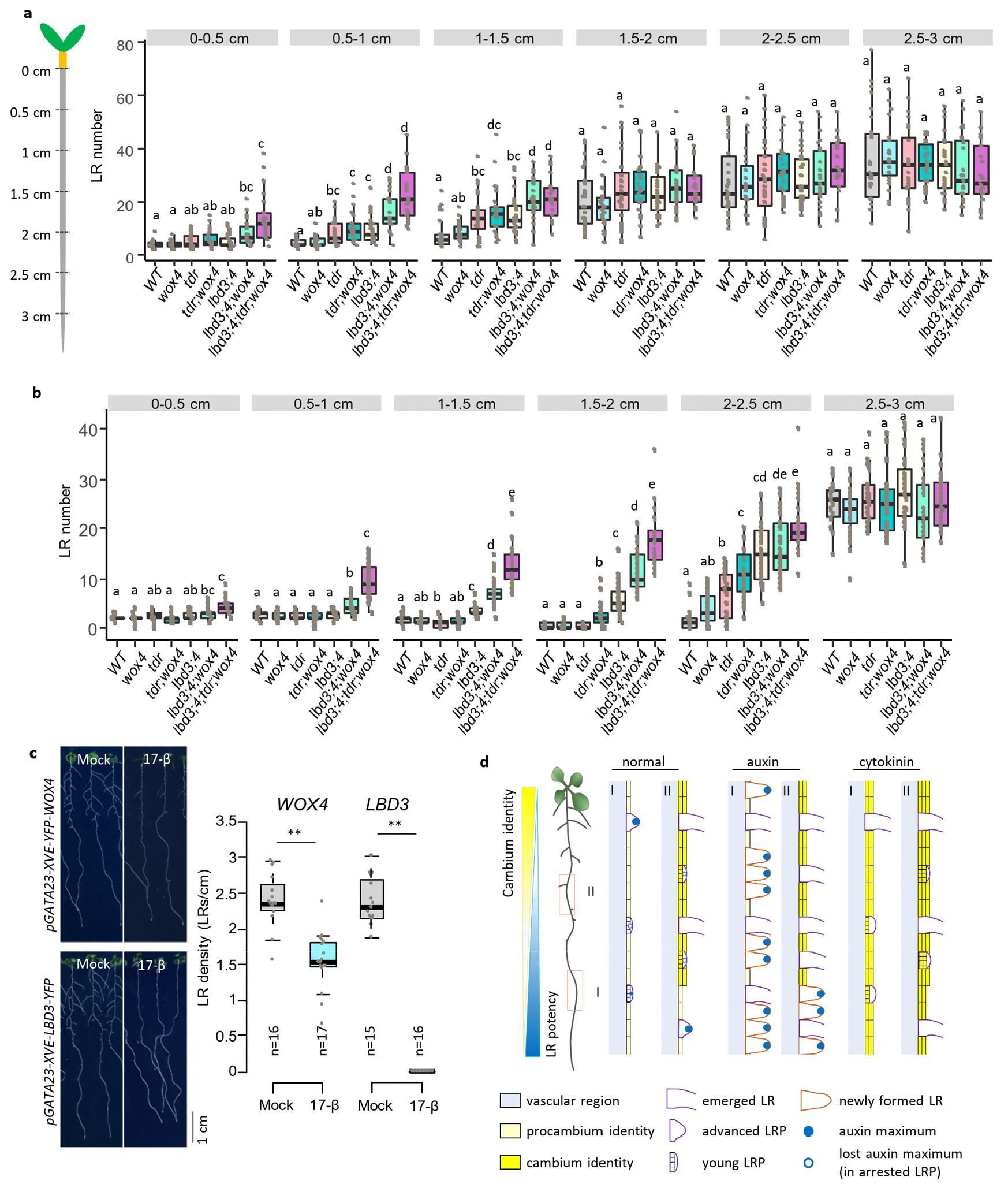
Cambium regulators shape the LR potency. **a-b**, Auxin-induced LR potency characterization of WT roots and cambium defective mutant roots. Seven-days old roots from distinct genotypes were treated 3-day with 5 pM NAA without (**a**) or with (**b**) a 2-day 1 pM BAP pretreatment. The left schematic (a) shows the strategy used to quantify emerged LRs in each root zone. Note *Ibd3;4* and *Ibd3;4;wox4* (**b**) exhibited a significant increase in LR induction in the root zone 1-1.5cm and 0.5-lcm, respectively, where a frequent cambium-like cell divisions occurred (**Supplementary Fig. 10**). A total number of 20-35 root samples were used for quantification in each genotype. The raw data are available in the source data file. Significant differences, indicated by different letters, were determined at an alpha level of 0.05 using a one-way ANOVA with either Tukey’s post hoc test (for equal homogeneous variance) orTamhane’s post-test (for unequal variance). Exact *p*-values for each comparison are provided in the source data file, **c**, Key cambium regulators are sufficient to inhibit LR development. The emerged LRs were quantified in each XPP-specific overexpression lines after 8-days germination on mock or induction medium, n, number of independent roots analyzed. Two-tailed Welch’s t test was performed. **, *p* < 0.01. The raw data are available in the source data file. Individual data points are plotted as grey dots (**a-c**). **d**, A simplified model illustrating how auxin and cytokinin balance cambium and LR fates of XPP cells.

We then analyzed the expression dynamics of cambium regulator genes within XPP cells and their derivatives in 7-day-old roots. Our analysis revealed a gradient expression pattern of cambium regulators within XPP cells and their derivatives, increasing towards the shoot, except for *PXY/TDR* (Supplementary Fig. 7a), which displayed an opposite trend (Supplementary Fig. 7b). Reporter analysis indicated that cambium-fate XPP cells gradually acquire cambium identity once they exit the root apical meristem, similar to the developmentally arrested LRPs (Figs. 2f and 3, and Supplementary Fig. 4a). This pre-established procambium identity can be conversely reverted to LR identity upon high levels of auxin treatment. For example, the expression of cambium regulator genes were clearly visible within the region 1.5-2 cm below the hypocotyl, where cambium was not yet active and massive LR induction was observed after auxin treatment (Supplementary Figs. 5b, 6 and 7a). However, when sustained expression of cambium regulators finally cause cambium activation, the reinforced cambium identity appears to block LR formation (Supplementary Figs. 5b, 6 and 7a).

To further explore the effects of cambium regulators on LR potency, we investigated cambium mutants and examined the relationship between their cambium activation and LR potency. For this purpose, we focused on mutants related to *PXY/TDR, WOX4, LBD3* and *LBD4*, which are known critical cambium regulators^38^ and loss-of-function mutant of them show a gradient of cambium activation defects from single, double, triple to quadruple mutants, as indicated from serial cross-sections to these mutants (Supplementary Fig. 6). While the mutants showed similar root length as WT and LR distribution after mock treatment (Supplementary Fig. 8), we again discovered a strong correlation between cambium activation defects and LR potency along the root after auxin treatment (Fig. 4a). For instance, the most severe mutant combination, *lbd3;4;tdr;wox4*, which exhibited no cambium activation in 7-day-old roots, showed efficient LR induction along the entire root length (Fig. 4a). Therefore, cambium regulators shape the LR potency along the root.

Next, we asked whether any of the cambial regulators are sufficient to prevent LR formation. We induced ectopic over-expressions of cambium regulator genes in the early LRPs and basal root region by using the *GATA23* inducible promoter. For this purpose, we included PXY/TDR, WOX4, LBD3, LBD4, LBD11, ANT and the HD-ZIP III transcription factor ATHB8, which acts downstream of auxin signaling in determining vascular stem cell organizer^5^. Ectopic expression analysis revealed that all these tested cambium regulators inhibited LR development to various degrees (Fig. 4c and Supplementary Fig. 9). PXY/TDR expression alone had no obvious effects on LR development, however, when combined with the treatment of its ligand, TRACHEARY ELEMENT DIFFERENTIATION INHIBITORY FACTOR (TDIF), a clear inhibitory effect was observed (Supplementary Fig. 9). Over-expressions of WOX4, LBD3 and LBD4 appeared to show stronger LR development inhibition compared with other cambium regulators, in line with broader LR potency in their corresponding mutants. In summary, the inhibitory effects of cambium regulators on LR development explain the restriction of LR potency exerted by cambium activation within XPP cells.

Overall, our results indicated that the pre-established procambium identity within XPP cells can be reverted to LR identity stimulated by high levels of auxin, further highlighting the cellular plasticity of XPP cells. However, gradual elevation of cambium regulator expression within XPP cells, especially TDIF-PXY/TDR, WOX4, LBD3 and LBD4, will lead to cambium activation and identity reinforcement, thereby serving as a limiting factor for LR potency.

### Cytokinins decreasing LR potency require cambium regulators

The previous findings have shown that pretreatment with cytokinin or expressing a cytokinin biosynthesis gene in the XPP cells diminishes the effectiveness of auxin induced LR formation, with the underlying mechanism unknown^12,13^. Given that cytokinins prompt premature cambium activation^19^ and they promote LR to cambium identity transition (Fig. 3), we formulated a hypothesis that cytokinins impede auxin-induced LR potency by facilitating cambium activities within XPP cells. To investigate this hypothesis, we conducted serial sections in the WT and cambium mutant roots following cytokinin treatment and quantified cambium activation events within XPP cells and their derivatives (Supplementary Fig. 10). Within the 0-2.4 cm root region below the hypocotyl, activation of cambium in XPP cells was evident in the WT root, which again correlated with the limited LR induction by auxin within this region (Fig. 4b, and Supplementary Fig. 10). Notably, cambium mutants exhibited an expanded auxin-induced LR potency subsequent to cytokinin pretreatment, with the *lbd3;4;tdr;wox4* mutant displaying the most pronounced LR potency (Fig. 4b and Supplementary Fig. 10). Interestingly, different from the WT, *lbd3;4* and *lbd3;4;wox4* demonstrated a significant increase in LR induction even from the root region where a frequent cambium-like cell divisions occurred (Fig. 4b and Supplementary Fig. 10). This finding suggests that cell divisions themselves within activated cambium may not serve as a limiting factor restricting LR potency, but the reinforced cambium identity, rendered by key cambium regulators, does so. Together, our results indicated that cytokinins play their inhibitory roles on auxin-mediated LR potency via promoting cambium activity. During this process, key cambium regulators are involved.

## Discussion

The developmental plasticity of plants ensures their ability to sustain growth in response to fluctuating environments. In this study, we delved into the fate plasticity of XPP cells. First, we developed novel LR-tracing systems, iRootTracker and RootTracker, to enhance LR/LRP visualization and demonstrated their broad applicability in various conditions. In contrast to cambium-fate XPP cells, developing LRPs exhibit distinct morphological features characterized by short cell walls and a domed shape, attributed to patterned cell divisions and differentiation. Traditional quantification of LR/LRP rely heavily on standard wide-field microscopes^39^, which could lead to the oversight of LRPs in root zones undergoing secondary growth or positioned non-orthogonally to the light path. To address this limitation, researchers often use LR reporters, albeit with drawbacks such as nonspecific or transient, stage-specific expression and faint expression in arrested LRPs. Our newly developed systems, which specifically, strongly and permanently marks LRPs post recombination, enable direct visualization of LRs/LRPs in intact roots under a fluorescent stereo microscope, therefore facilitating convenient and high-throughput LR-related quantification. In addition to the 35S-Loxp-erGUSpYFP construct used in this study, we also provided five more vectors for the research community by replacing 35S promoter with other two frequently used constitutive promoter UBIQUITIN10^40^ and HTR5^41^, or by changing the coding sequence of YFP to RFP.

Using the root iRootTracker and cambium reporters, we revealed that approximately half of LRPs in the mature root zone became arrested and eventually acquired cambium identity under our growth conditions (Figs. 2 and 4d). Furthermore, we demonstrated that cytokinins promoted arrest of young LRPs and the transition from LR identity to cambium identity, consistent with their essential roles in secondary growth^18,19^ (Figs. 2b, c,3 and 4d). A well-established auxin maximum is believed to play a crucial role in the development of LRP^15,35^. Our results suggest that LRP arrest might result from a failure to maintain this auxin maximum. This was supported by our observation that supplementing low concentrations of auxin to a 4-day-old root was effective in stimulating the emergence of most LRPs, while higher concentrations of auxin could awaken arrested LRPs in a 7-day-old root (Figs. 2b, c and 4d and Supplementary Fig. 3c). However, as the root matures, the capacity of auxin to reactivate arrested LRPs diminishes (Fig. 4a, d and Supplementary Fig. 5a, b), because these LRPs have already acquired and reinforced their cambium identity (Figs. 2f and 4d).

In this study, expression analysis of cambium regulators revealed a gradual acquisition of cambium identity of cambium-fate XPP cells, as root matures (Fig. 4d and Supplementary Fig. 7a). When expression of cambium regulator genes reach a threshold level, cambium activation occurs. Even though in this study only XPP cells and their derivatives were followed, it is likely that in the vascular region, procambium activation to cambium occurs in a similar manner. With the RootTracker, we showed that the pre-established procambium identity within XPP cells could be reverted to LR identity by auxin treatment (Fig. 4a, d, and Supplementary Figs. 5b and 7). When cambium activation occurs, the XPP derivatives become less responsive to auxin-induced LR formation, likely due to their strengthened cambium identity, boosted by the key cambium regulators PXY/TDR, WOX4, LBD3 and LBD4. Supporting this, overexpression of these regulators strongly inhibited LR development and the corresponding knockouts displayed expanded LR potency (Fig. 4c and Supplementary Fig. 9). It is possible that other cambium regulators also contribute to this process, such as ATHB8, ectopic expression of which showed strong LR development inhibition (Supplementary Fig. 9).

It has been shown that auxin cannot rescue the LR initiation defect when expressing an IPT gene in XPP cells or giving cytokinin pretreatment^12,13^. In this study, our data showed that cytokinin pretreatment lowers LR potency via premature cambium activation within XPP cells in a process requiring at least PXY/TDR, WOX4, LBD3 and LBD4 (Fig. 4b, d and Supplementary Fig. 10). This represents another pathway blocking LR development, which is likely different from their direct inhibitory effects on LR development via modulating PIN1-mediated auxin transport^14^. A prior study indicated that while LR formation is the predominant response to auxin treatment in 4-day-old seedlings, for 6-day-old seedlings, auxin treatment induces periderm formation instead, in the mature part of the root^34^. This phenomenon has been observed earlier as well^36^. This dual role of auxin can potentially be explained by our findings. Our data suggest that once accumulation of cambium regulators promotes periderm identity in XPP and the remaining pericycle cells, then auxin has a new role to promote further periderm development. In conclusion, our findings demonstrated the control of cell fate plasticity of XPP cells in *Arabidopsis* roots and elucidated how plant hormones auxin and cytokinin influence root architecture and secondary growth through balancing LR fate and cambium cell fate of XPP cells.

## Methods

### Vector construction and *Arabidopsis* transformation

To generate the *pXPP-XVE* inducible promoter, a 2.5-kb XPP-specific promoter^37^ was cloned into the first box entry vector *1R4a-ML-XVE*^42^ by restriction enzyme digestion and ligation. To facilitate the construction of inducible promoters, we modified the *1R4a-35S-XVE* vector (“a” denotes ampicillin resistance) by adding Asc I and Fse I restriction enzyme sites flanking the *35S* promoter sequence through omega PCR^43,44^, yielding the *1R4a-AscI-35S-FseI-XVE* vector. Subsequently, a 1.1-kb *GATA23* promoter sequence was introduced by replacing the *35S* promoter via digestion and ligation. The *1R4a-35S-XVE* vector was further modified by incorporating multiple cloning sites flanking the *35S* promoter using omega PCR^43,44^ to generate a universal intermediate vector, *1R4a-MCS-35S-MCS-XVE*. A 2-kb *HB53* promoter sequence was first integrated into the first box donor vector pDONR1R4z (“z” denotes zeocin resistance) via BP reaction, to produce the *1R4z-pHB53* entry vector. The HB53 promoter sequence was also cloned into the *1R4a-MCS-35S-MCS-XVE* vector through digestion and ligation to produce *1Ra-pHB53-XVE*. The RFP coding sequence from the published *221z-erGUSpRFP* entry vector (“z” denotes zeocin resistance) ^45^ was replaced with YFP coding sequence via omega PCR^43,44^, to generate *221z-erGUSpYFP*.

A longer version of *35S* promoter^46^ and a 2-kb *HTR5* promoter^41^ were used to substitute for the shorter version of *35S* promoter in the published *1R4z-35S-Loxp* entry vector^5^ through omega PCR^43,44^, for the generation of *1R4z-p35S*_*long*_*-Loxp* and *1R4z-pHTR5-Loxp*, respectively. To avoid misunderstanding, the *35S*_*long*_*-Loxp* in this paper is presented as *35S-Loxp*. The construction of the *1R4z-pUBQ10-loxp* vector was performed with a different strategy. Initially, multiple cloning sites were introduced into the flanking sites of the *35S* promoter of the published *1R4z-35S-Loxp* entry vector^5^ via omega PCR^43,44^ to produce a common intermediate vector *1R4z-MCS-35S-MCS-loxp* entry vector, which was followed by the replacement of the *35S* promoter with a 2-kb *UBQ10* promoter sequence through digestion and ligation.

All binary expression constructs were generated through multisite gateway LR reactions. The promoters *1R4a-GATA23-XVE, 1R4a-HB53-XVE* and *1R4z-HB53* in the first boxes, recombinase coding sequences 221z-CRE^5^ and 221a-CYCB1;1-CRE in the second boxes^5^, and the terminator *2R3a-nosT* in the third box were combined into the destination vector FRm43GW^44^. The first box entry vectors *1R4z-35S-Loxp, 1R4z-pHTR5-Loxp* and *1R4z-pUBQ10-Loxp*, second box entry vectors *221z-erGUSpYFP* and *221z-erGUSpRFP*^45^, and the third box entry vector *2R3a-nosT* were integrated into the destination vector *pBM43GW*^47^. In this study, we tested the *35S-Loxp-erGUSpYFP* construct by first transforming it into Col-0 background. Then in T2 generation, 15 independent, likely single insertion lines (as indicated by Mendelian segregation ratio) were screened and transformed with *pGATA23-XVE-CRE*. The most suitable *35S-loxp-erGUSpYFP* line was selected based on performance (i.e. consistent and non-leaky clone formation) with *pGATA23-XVE-CRE*, and that line (without *pGATA23-XVE-CRE*) was used as the background for subsequent studies.

The reporter lines used in this study, *DR5erRFP*^42^, *pPXYerYFP*^5^, *pWOX4erYFP*^5^, *pLBD3erYFP*^19^, *pLBD4erYFP*^19^, *pANTerRFP*^45^, *pWOX5erGFP*^48^, *gPLT1YFP*^49^ have been published before. A 2-kb promoter sequence of PER15^34^ was cloned into the first box donor vector *1R4z-BsaI-ccdB-BsaI*^45^ via goldengate cloning to produce *1R4z-PER15*. Entry vectors *1R4z-PER15, 221z-erYFP*^*42*^ and *2R3a-nosT*^*42*^ were combined into the destination vector *pHm43GW*^*47*^ to generate the pPER15erYFP reporter construct.

Constructions of *221z-ANT, 221z-ATHB8* entry vectors were described elsewhere^5^. The coding sequences of *LBD3, LBD4*, and *LBD11* without stop codons and the genomic sequence of *PXY/TDR* with the stop codon were cloned into the second box donor vector *221z-BsaI-ccdB-BsaI*^44^ through goldengate cloning. The coding sequences of *YFP* and *WOX4* were initially separately amplified in the first round PCR and then combined into a fusion of *YFP-WOX4* through a second round overlapping PCR, which was followed by a subsequent BP reaction to produce the *221z-YFP-WOX4* entry vector. For cambium regulator ectopic expression analysis, the inducible promoter *1R4a-pGATA23-XVE*, respective cambium regulator genes, and *2R3a-nosT* (for *PXY/TDR* and *YFP-WOX4*) or *2R3a-YFP*^42^ (for other cambium regulator genes) were integrated into either *pBm43GW*^47^ or *pFRm43GW*^44^ destination vectors. All constructs were transformed into Arabidopsis Col-0 background. For each construction, more than 10 lines were analyzed and one representative line showing consistent expression or phenotype with others was used for further analysis. All primers used in this study are listed in the Supplementary Table 1. All constructs generated in this study are listed in the Supplementary Table 2.

The cambium defective mutants used in this study *wox4, tdr, tdr;wox4, lbd3;4, lbd3;4;wox4* were published elsewhere^19^. The mutant combination *lbd3;4;tdr;wox4* was produced by crossing *lbd3;4* with *tdr;wox4*.

### Plant growth and chemical treatment

Seeds were surface sterilized and stratified for 2 days at 4 °C before plating on half-strength MS growth medium containing 0.5 × MS salt mixture with vitamins (Duchefa), 1% sucrose, 0,5g/l MES pH 5.8 and 0.8% agar. Transgenic seeds were screened either under a fluorescent stereo microscope (Leica M165 FC) or on growth medium containing 20 /ml phosphinotricin (Sigma) or 20 /ml hygromycin (Sigma). Plates were vertically positioned in the growth chamber at 23°C with long-day settings (16h light and 8h dark). Stock solutions of 10mM 1-Naphthylacetic acid (NAA, Sigma) and 10mM 6-benzylaminopurine (BAP; Sigma) were prepared in dimethyl sulfoxide (DMSO, Sigma) and diluted into required working concentrations. 20mM 17-

-estradiol (17-β, Sigma) was dissolved in DMSO as stock solution, and diluted to 5 M as working concentration. Potassium Nitrate (Sigma), mannitol (Sigma), and sodium chloride (Sigma) were added to the growth medium to indicated concentrations. Auxin treatment in LR-tracing experiments was performed on solid growth medium with 0.8% plant agar. For LR induction experiments performed in WT and cambium defective mutants, liquid growth medium with 0.2% plant agar was used. An equal volume of DMSO solution was added to growth medium as mock treatment.

### GUS staining and histological sections

GUS staining were performed as previously described^5^. Serial longitudinal sections and serial cross sections were carried out as described previously^50^. For vibratome sections, 7-day-old root samples were first fixed in 4% paraformaldehyde solution (in 1x PBS, pH 7.2, Sigma) overnight, then rinsed twice with 1x PBS, aligned manually on a glass slide placed on ice, and cut every 0.5cm. The segmented root samples were then embedded in 4% agar (Sigma, dissolved in 1x PBS) for subsequent vibrotome sections. Vibratome sections was carried out as described before^5^. Root sections were stained in 1x PBS solution containing 0.1% calcofluor for cell wall staining.

### Microscopy and image processing

Intact LRs/LRPs observation and tracing in a time-course manner was performed under the fluorescence stereo microscope (Leica M165 FC). GUS-stained root samples were first fixed into 4% PFA solution overnight, followed by an incubation in ClearSee solution^51^ for at least 2days before LRs/LRPs observation and quantification under a standard wild filed microscope (Leica 2500). Confocol microscopy (Leica SP8) of the lateral view of LRs/LRPs involves fixing root samples in a 4% paraformaldehyde solution overnight, followed by incubation in ClearSee for at least two days, and 1-day cell wall staining with 0.1% calcofluor dissolved in ClearSee. This method was also used for lateral view observation of reporter expressions in LRPs. The cross sections of WT and cambium defective mutants were stained sequentially in a 0.05% (w/v) ruthenium red solution (Fluka Biochemika) and a 0.05% (w/v) toluidine blue solution^5^ in deionized water before observation. The figures were organized in PowerPoint. Images for LR tracing sometimes were adjusted in brightness and contrast for better visualization of LRPs with weak signals. Images used for comparison were always captured and displayed with the same settings.

### Quantifications and Statistics

Root length quantification in 7-day-old roots were performed by first scanning the plates where the seedlings were grown then measuring root length in scanned images with Fiji-ImageJ. Emerged LR numbers of WT roots and cambium defective mutant roots after a 3-day auxin or mock treatment were quantified by first subjecting the seedlings to a 70°C oven for two hours to prevent further growth, then placing roots on a glass slide with a ruler underneath and counting root numbers every 0.5 cm. The numbers of emerged LRs in ectopic overexpression lines of cambium regulators were counted under the stereo microscope, after an 8-day germination on mock or induction growth medium. Fluorescence intensity of different cambium reporters within XPP cells/derivatives was quantified using Fiji-ImageJ.

Plots were created using the boxplot function and the ggplot2 package in RStudio (https://www.rstudio.com/) with R version 4.3.3 (https://www.r-project.org/). The boxplots show the first quartile (bottom of the boxes), median (middle line), and third quartile (top of the boxes). Individual samples are represented by dots. All experiments were conducted at least twice. Statistical analyses were performed with SPSS Statistics version 29 and RStudio (https://www.rstudio.com/) using R version 4.3.3 (https://www.r-project.org/). For comparing two groups, the Welch’s t-test was applied following the performance of a Shapiro test. For multiple group comparisons, one-way ANOVA was conducted in SPSS Statistics version 29. Levene’s Test assessed the homogeneity of variances. Significant differences between datasets were determined using either the Tukey post hoc test (for equal homogeneous variance) or Tamhane’s post-test (for unequal variance) at a significance level of alpha = 0.05. All ANOVA results are detailed in source data files.

## DATA AND MATERIALS AVAILABILITY

All vectors and plant material in this study are available from the corresponding authors Xin Wang and Ari Pekka Mähönen upon request.

## ACKNOWLEDGEMENT

We thank J. López Ortiz, X. Zhang, H. lida, M. LV and T. Blomster for providing feedback on the manuscript. We thank H. Mulvey for help cloning the XPP inducible promoter. Confocal imaging was supported by the Light Microscopy Unit (LMU), University of Helsinki. This work was supported by the Academy of Finland (grant numbers 316544 and 346141 to A.P.M.); European Research Council (ERC-CoG CORKtheCAMBIA, agreement 819422 to X.W., L.Y. and A.P.M.); Wallenberg Academy Fellowship from the Knut and Alice Wallenberg Foundation (to C.W.M.). X.W. was also supported by a grant from the Chinese Scholarship Council (CSC). The authors also acknowledge the use of ChatGPT for wording improvement and take full responsibility for the content of the publication.

## CONTRIBUTIONS

X.W. and A.P.M designed the experiments and X.W. performed all experiments with the help of L.L.Y.

L.L.Y. performed statistical analysis. J.Z. and C.W.M. provided unpublished material. X.W. and A.P.M wrote the paper with input from all authors.

