## Supplementary Material for "Cell fate plasticity of xylem-pole-pericycle in *Arabidopsis* roots"

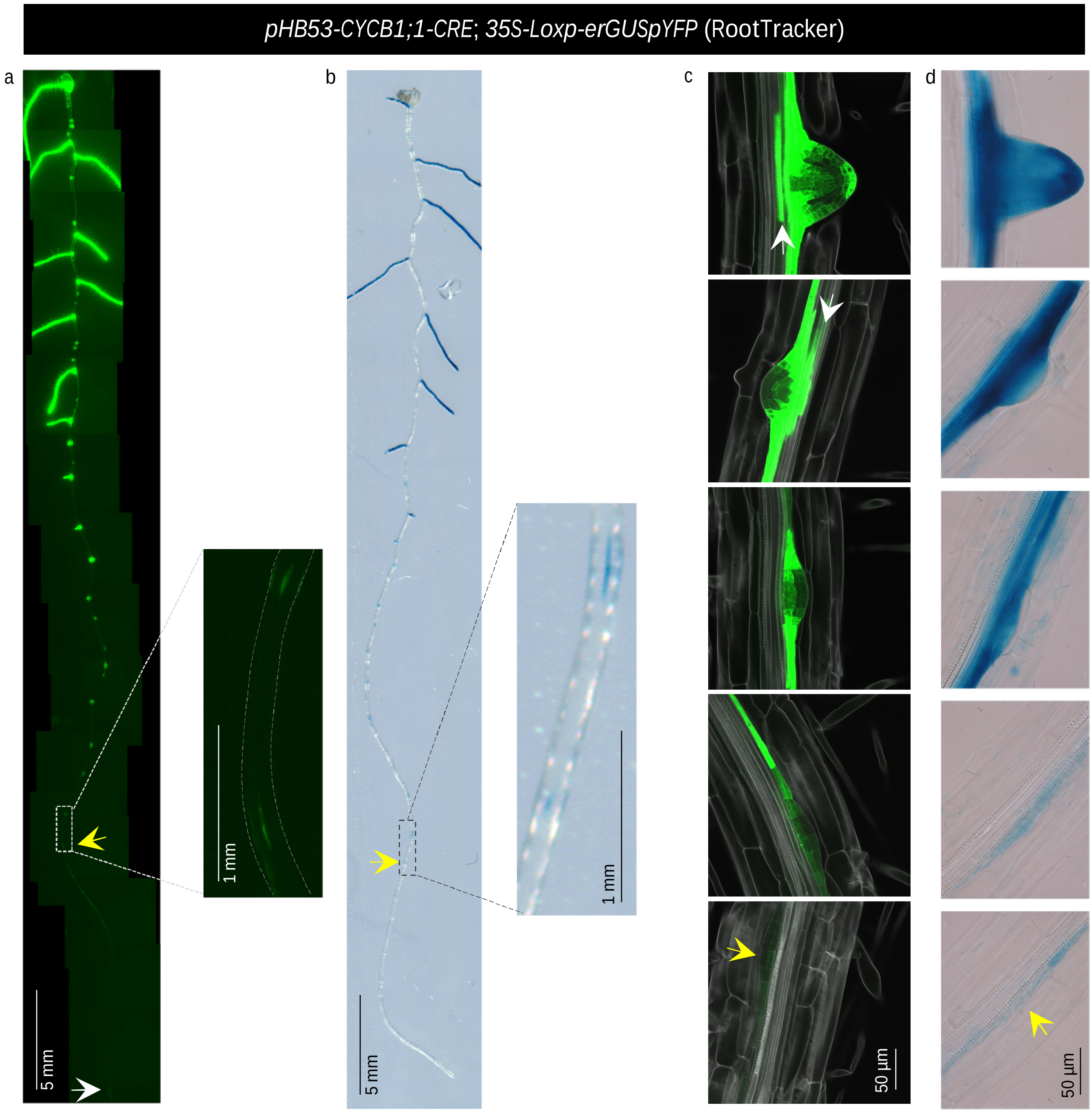


Supplementary Fig. 1: Generation of a stable LR-tracing system (RootTracker) with the *pHB53* native promoter.

a-b, An example showing LR/LRP labeling with YFP (a) or GUS (b) in 7-day-old roots under the fluorescent stereo microscope using the YFP channel (a) or the bright field (b). Framed root region with weak YFP (a) or GUS (b) marking is magnified in respective right panel. c, A careful confocal inspection showed YFP clones faithfully mark LRs/LRPs (395/395, n=12), and LRs/LRPs were effectively labeled by YFP signals (216/220, n=6). White arrows indicate the unspecific YFP signals associated with overlying LRPs. d, Standard microscopy observation post-GUS staining revealed GUS clones precisely predict LRs/LRPs (365/365, n=12) and LRs/LRPs were effectively labeled with GUS staining (365/378, n=12). White arrow marks the root tip (a). Yellow arrows show weak signals in young LRPs near the root tip (a-d).

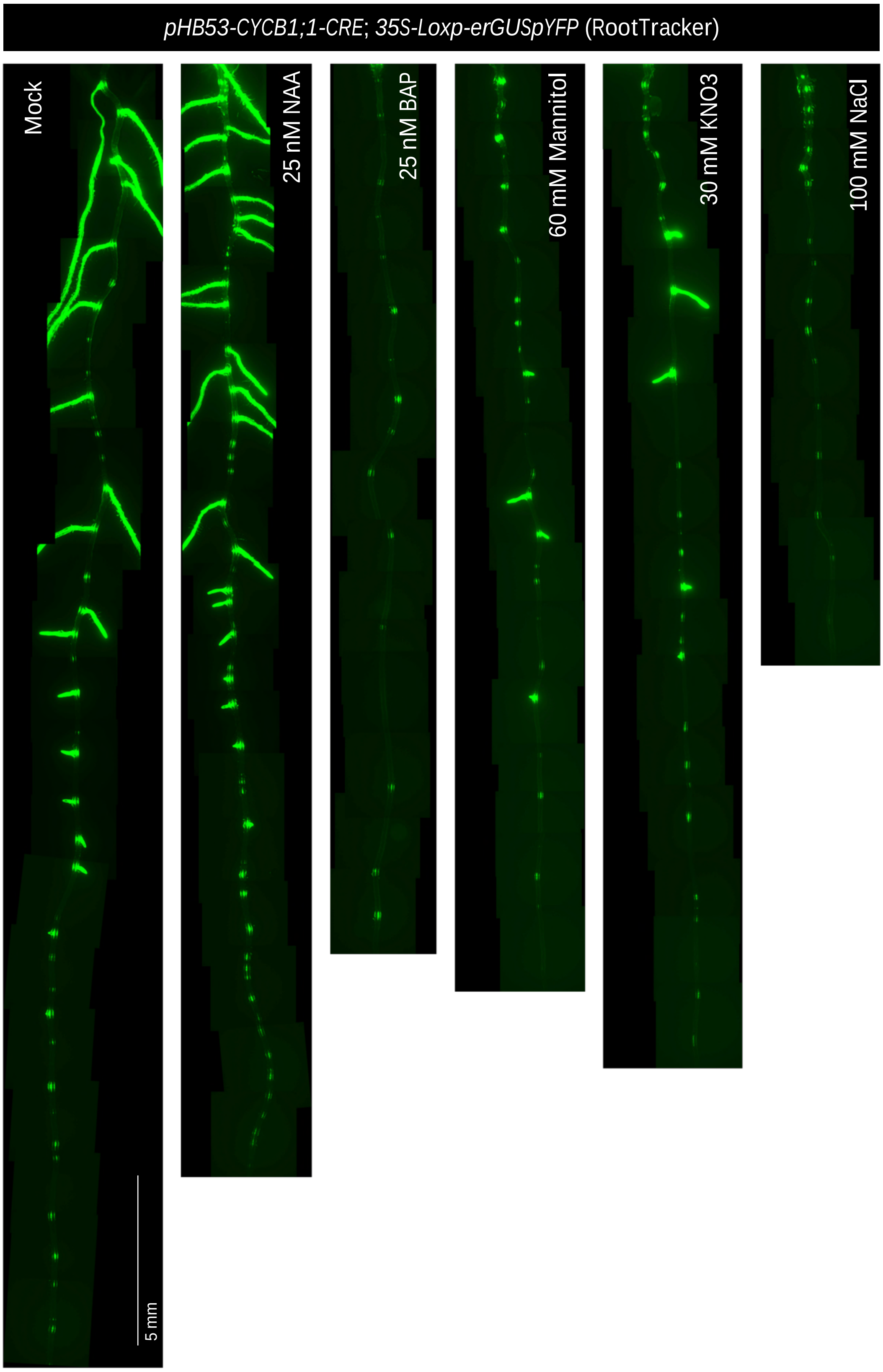


Supplementary Fig.2: Wide applicability of the RootTracker under various conditions. After 8-days germination of the RootTracker line on each indicated growth medium, the root architecture were visible under a fluorescent stereo microscope. While auxin promotes LR outgrowth (fluorescence in the whole, outgrown LR), cytokinins inhibit LR outgrowth and promote LRP arrest (dot-like green signal). Under drought (mannitol), high nitrate and salt conditions, suppression of LR outgrowth and increase in arrested LRPs was observed. For each condition, n>30.

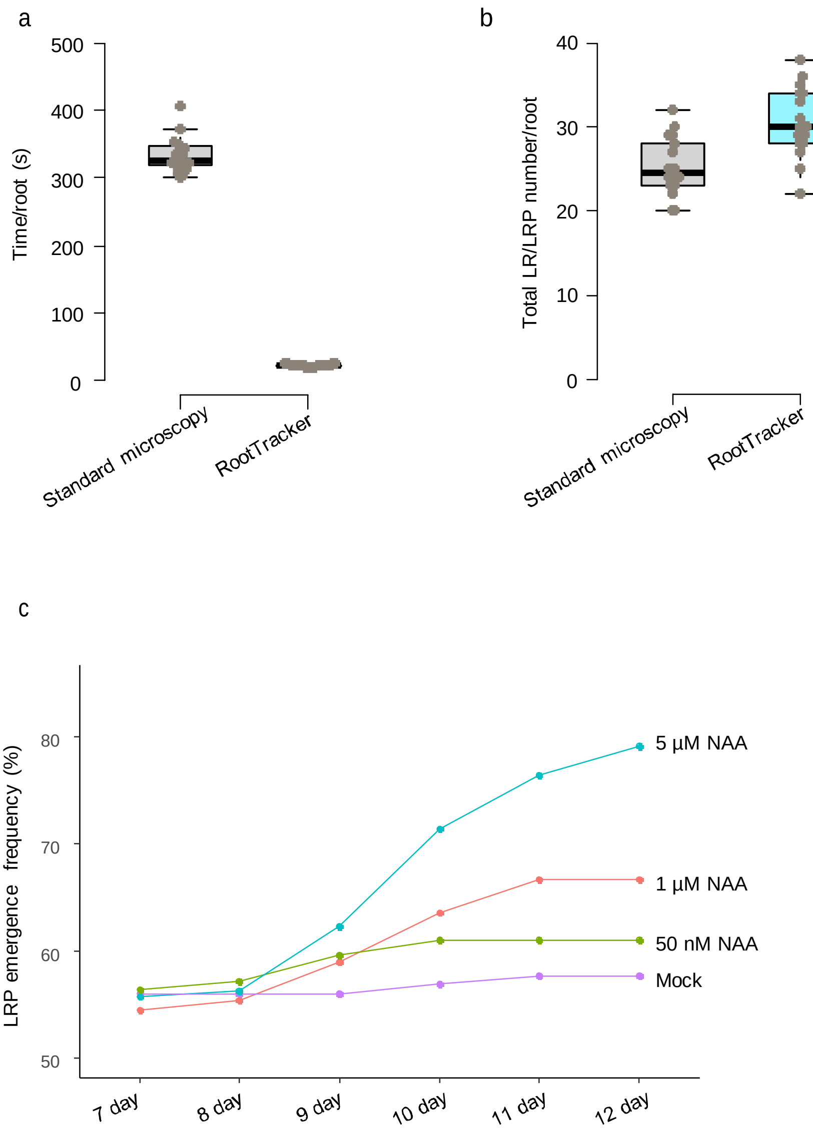


Supplementary Fig. 3: The novel LR-tracing systems enable fast and convenient LR analysis.

a-b, A comparison of analysis time on LR/LRP number quantification per root (a) and the number of quantified LR/LRPs per root (b) between the standard microscopy and the RootTracker. Seven-day-old roots from the RootTracker line were used. The same root was first analyzed under a fluorescent stereo microscope then fixed and cleared before standard microscopy analysis. Gray dots represent individual data point, with 18 roots used for quantification. The raw data are available in the source data file. c, LR/LRP tracing in 7-day-old roots with different supplied auxin levels by using the iRootTracker. The 3-days old seedlings were first induced for 1-day, then transferred to estradiol-free medium for another 3 days before tracking each LR/LRP. With 5 µM NAA treatment, we also observed 22 newly emerged LRs which were not labeled with YFP. The data are shown as mean values calculated from a total LRP number of 139 to 330, based on 15 to 42 individual roots. The raw data are available in the source data file.

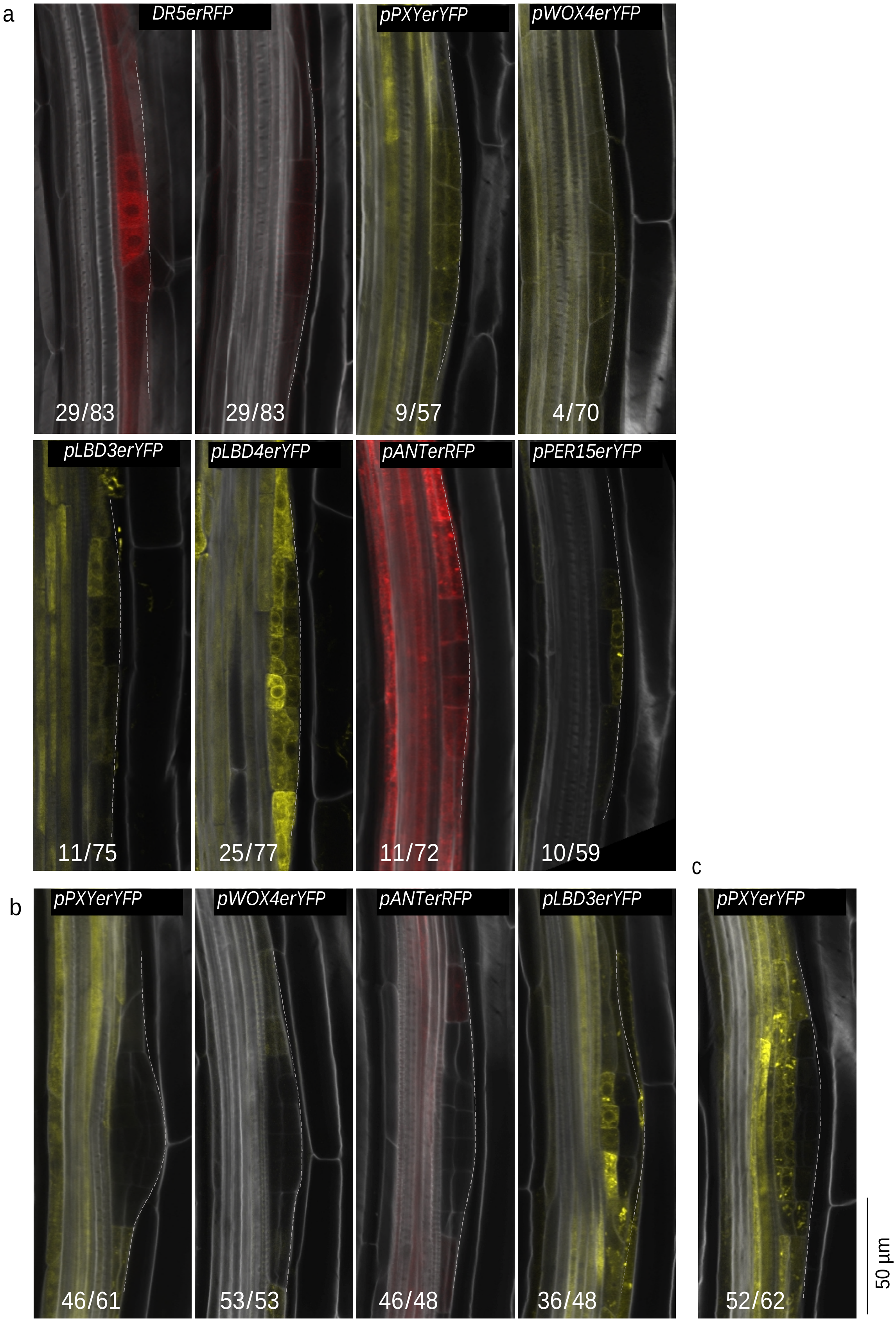


Supplementary Fig. 4: Arrested LRPs gradually obtain cambium identity and cytokinins promote this transition.

a, In 5-day-old roots with mock treatment, a subset of LRPs displayed low DR5 activity and expression of cambium reporters. A similar stage LRP with normal DR5 activity is shown in the left panel as control. Note these LRPs are all morphologically at young stage. Numbers indicate the observed frequencies of LRPs (out of total lateral organs) with cambium regulator gene expression. b, A 10h 1 µM BAP treatment to 3-days old roots was insufficient for cambium regulator gene expression in LRPs, except *LBD3*. For *PXY* expression in most LRPs, a 1-day induction was required (c). Numbers indicated the frequencies of observed LRPs with/without cambium regulator gene expression (b, c). LRPs are outlined with white dashed lines.

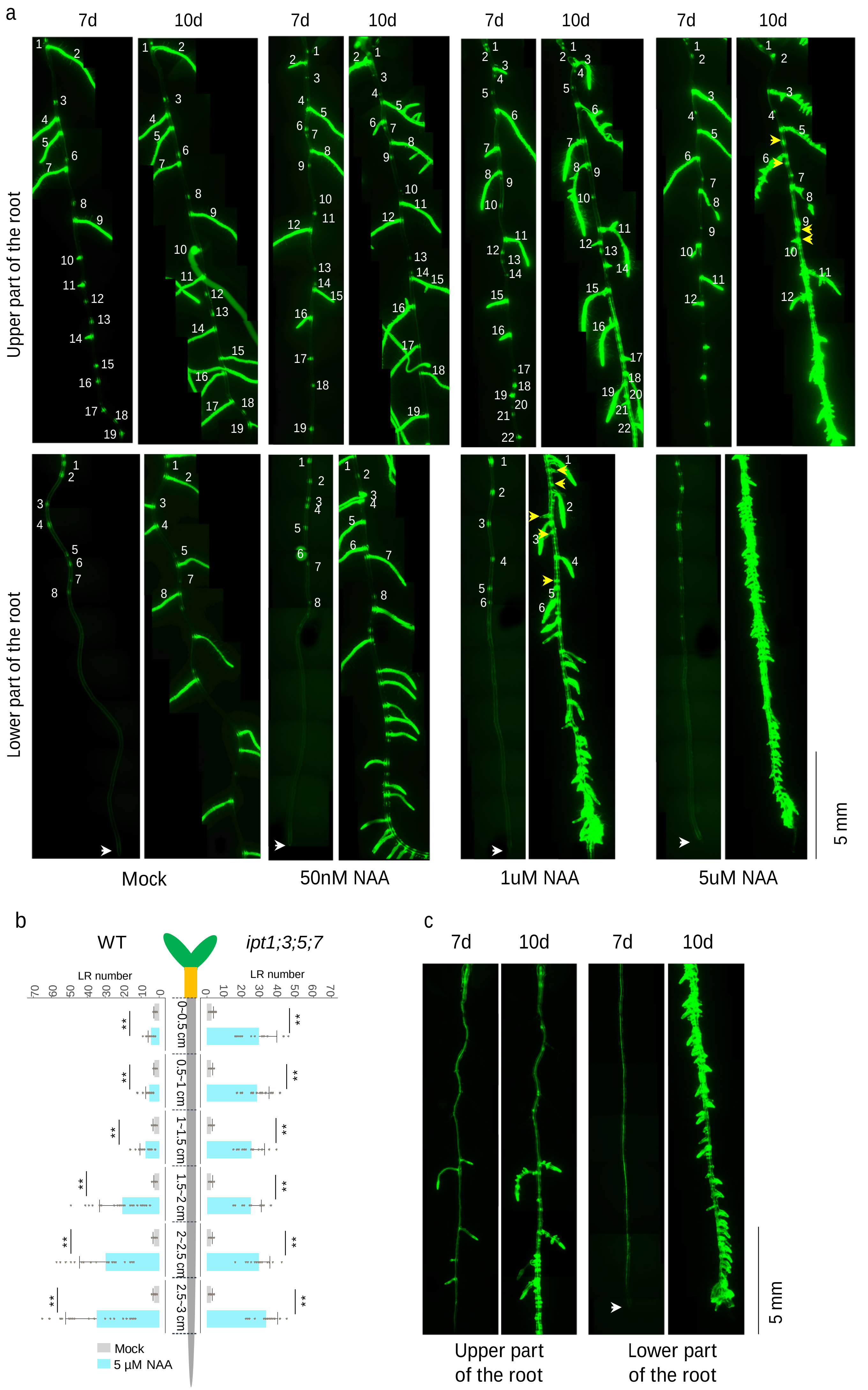

a
7d
10d
Upper part of the root
Lower part of the root
Mock
50nM NAA
1uM NAA
5uM NAA
5 mm
b
WT
ipt1;3;5;7
LR number
0~0.5 cm
0.5~1 cm
1~1.5 cm
1.5~2 cm
2~2.5 cm
2.5~3 cm
Mock
5 µM NAA
c
Upper part of the root
Lower part of the root

Supplementary Fig. 5: Auxin-induced LR-potency characterization.

a, Auxin-induced LR potency assay along the 7-day-old roots before (left) and after a 3-day auxin treatment (right), with the RootTracker. The same number marks the same LR/LPR before and after treatment. Only LRs/LRPs that can be distinguished from newly formed LRs/LRPs (yellow arrows) are marked. b, Quantifications of emerged LR numbers along the roots after a 3-day 5 μM NAA treatment to 7-day-old roots. Quantification was performed every 0.5 cm below the hypocotyl as the schematic indicated. The data are presented as the mean ± SE derived from observations collected from 15 to 34 individual roots. Two-tailed Welch's t test was performed. **, $p < 0.01$. The raw data are available in source data file. Note that massive LR induction occurs from the region 1.5-2 cm towards the root tip. In the upper 1 cm, only slightly increased number of LR was observed compared to control. Our LR-tracing result showed that these newly emerged LR originate at least partially from the reactivation of arrested LRPs (Supplementary Fig. 3c). 1-1.5 cm seems to be a transition zone, where more LR induction was observed than the above zone. c, Newly formed LRs/LRPs triggered by a 3-day 5 μM NAA treatment were originated from the XPP cells and their derivatives, as revealed from the XPP cell lineage tracing system (*pXPP-XVE-CYCB1;1; 35S-Loxp-erGUSpYFP*). To mark the XPP cells with this system, 4-days old seedlings were induced for 3-days before transfer to 5 μM NAA medium for another 3 days. For each treatment, n>20 (a, c). White arrows mark the primary root tips (a, c).

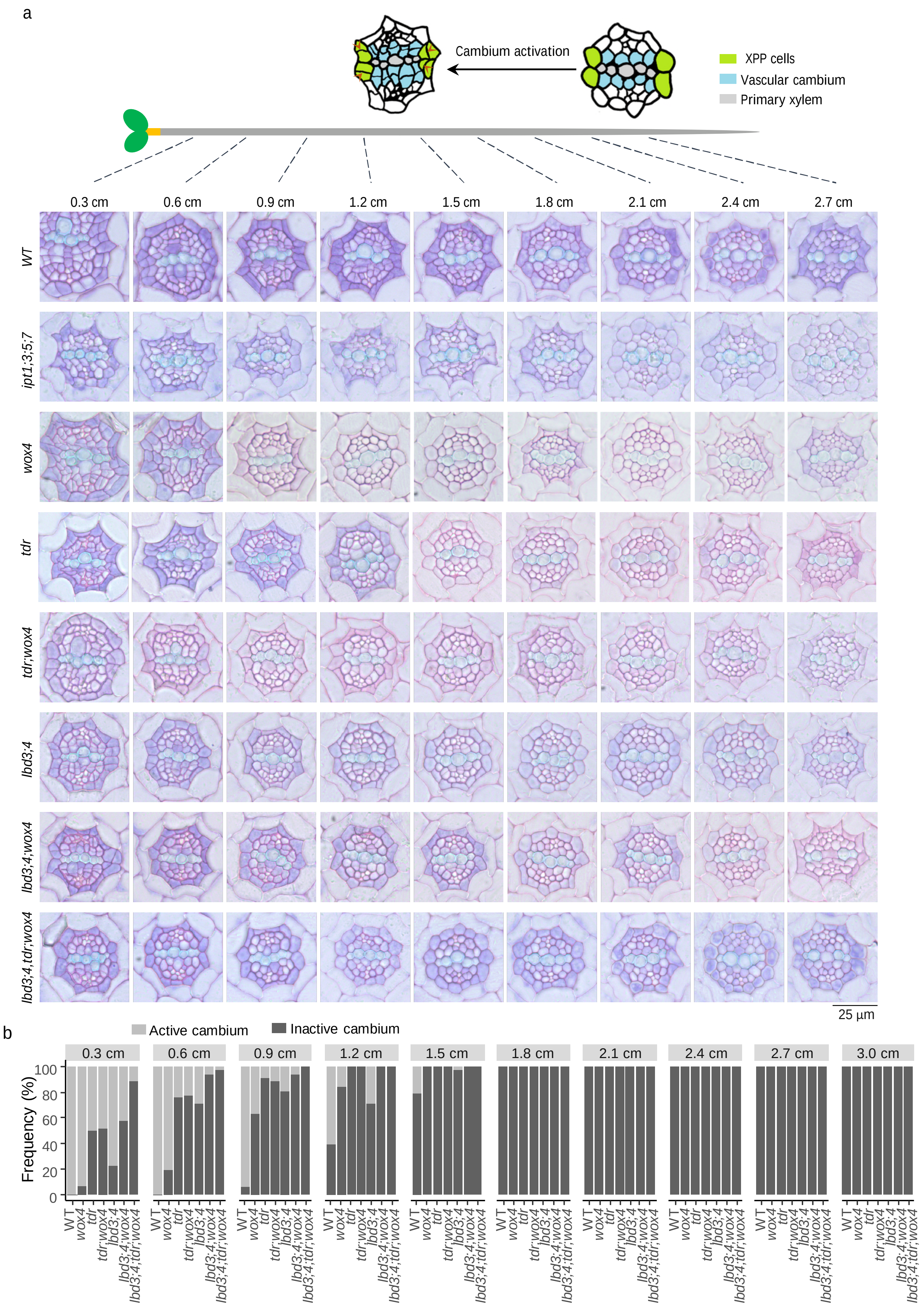


Supplementary Fig.6: Cambium activation assay within XPP cells of WT roots and cambium defective mutant roots. a, Serial cross sections of 7-day-old WT roots and cambium defective mutant roots. The schematic on the top: when cell divisions occur on both XPP poles, we considered cambium to be activated. b, Quantification of cambium activation frequency within XPP cells along the 7-day-old roots of different genotypes. A total of 31 to 35 individual roots were used for the quantification. The raw data are available in the source data file.

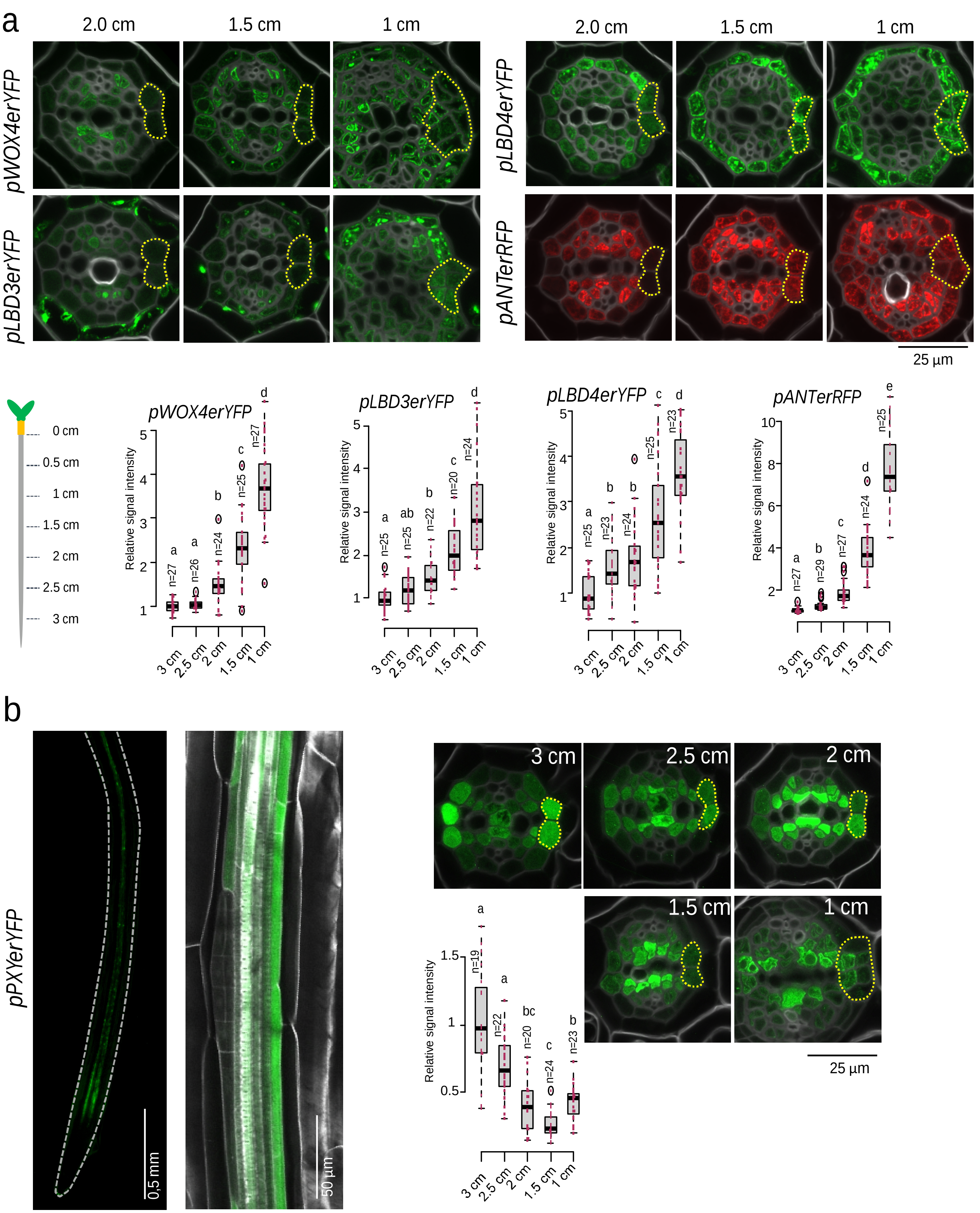


Supplementary Fig. 7: Gradual acquisition of cambium idenity in XPP cells.

a, Gene expression analysis of cambium regulators in 7-day-old roots revealed a gradient expression within XPP cells and their derivatives (marked by the dotted line) as the root matures. The schematic shows the relative position of each section along the root. b, *PXY/TDR* displays an opposite expression gradient with relatively strong and specific expression in XPP cells above the root tip region. Cell walls were visualized by calcofluor. Red dots indicate relative signal intensity in individual roots. The raw data are available in the source data file. Significant differences, indicated by different letters, were determined at an alpha level of 0.05 using a one-way ANOVA with either Tukey's post hoc test (for equal homogeneous variance) or Tamhane's post-test (for unequal variance). Exact *p*-values for each comparison are provided in the source data file.

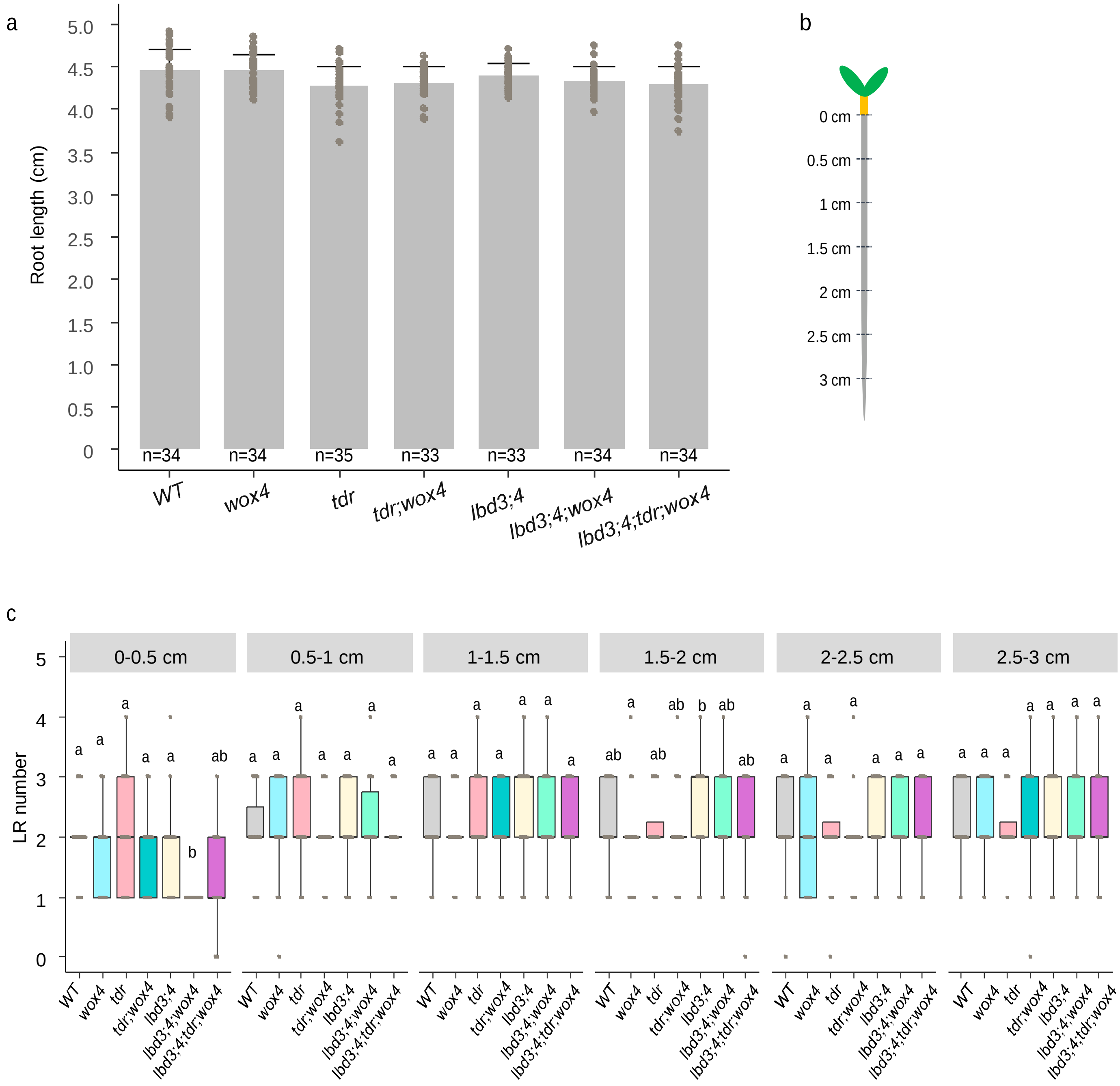


Supplementary Fig.8: Comparisons of root length and emerged LR distributions among WT and cambium defective mutants.

a, Similar root length observed among different genotypes in 7-day-old roots. b, A schematic shows the strategy used to quantify emerged LR numbers in each root zone (c) in 10-day-old roots with mock treatment. Quantification was performed using 25-35 individual roots, represented by gray dots on the boxplot. The raw data are available in the source data file. Significant differences, indicated by different letters, were determined at an alpha level of 0.05 using a one-way ANOVA with either Tukey's post hoc test (for equal homogeneous variance) or Tamhane's post-test (for unequal variance). Exact *p*-values for each comparison are provided in the source data file.

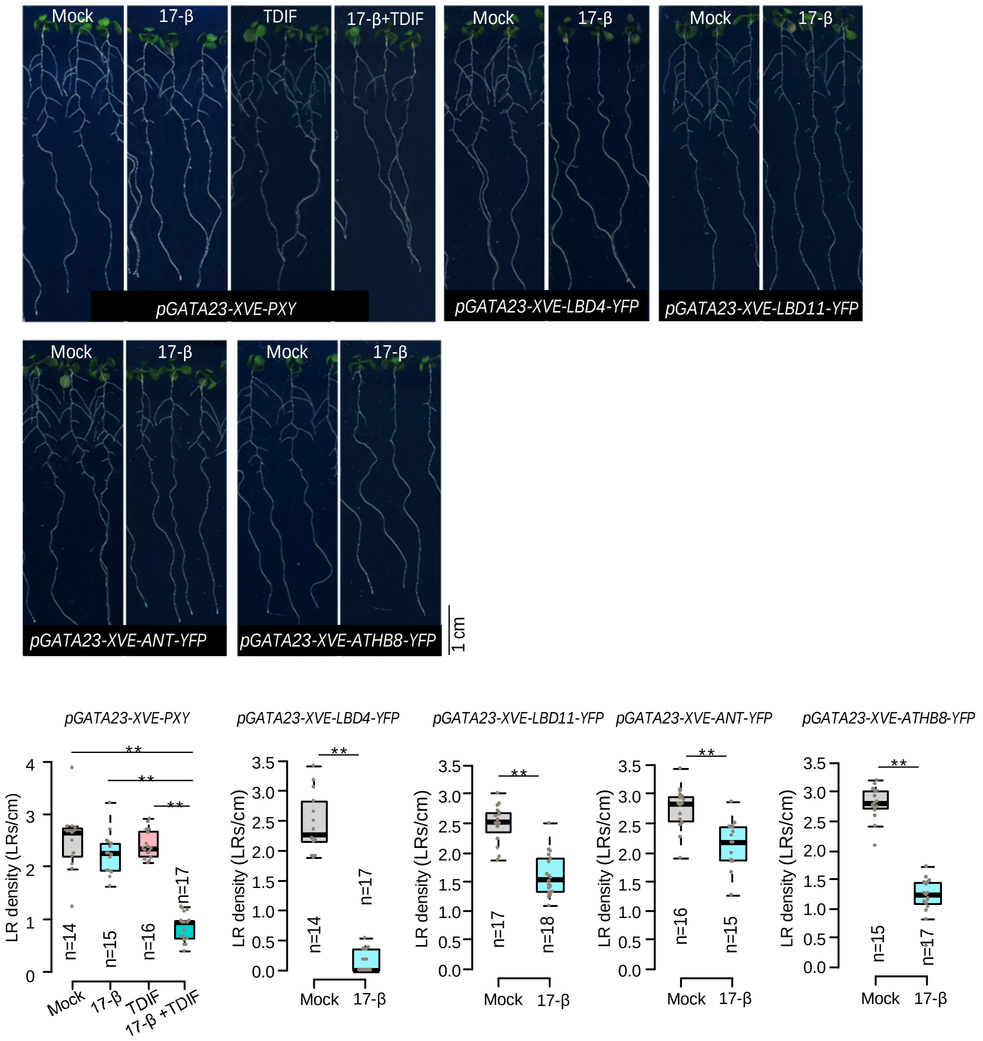


Supplementary Fig. 9: Ectopic expression of cambium regulators is sufficient to inhibit LR development. The emerged LR numbers were quantified under the stereo microscope after 8-days germination on mock/induction growth medium. Individual data points are plotted as grey dots. n, number of independent roots analyzed. Two-tailed Welch's t test was performed. **, $p < 0.01$. The raw data are available in the source data file.

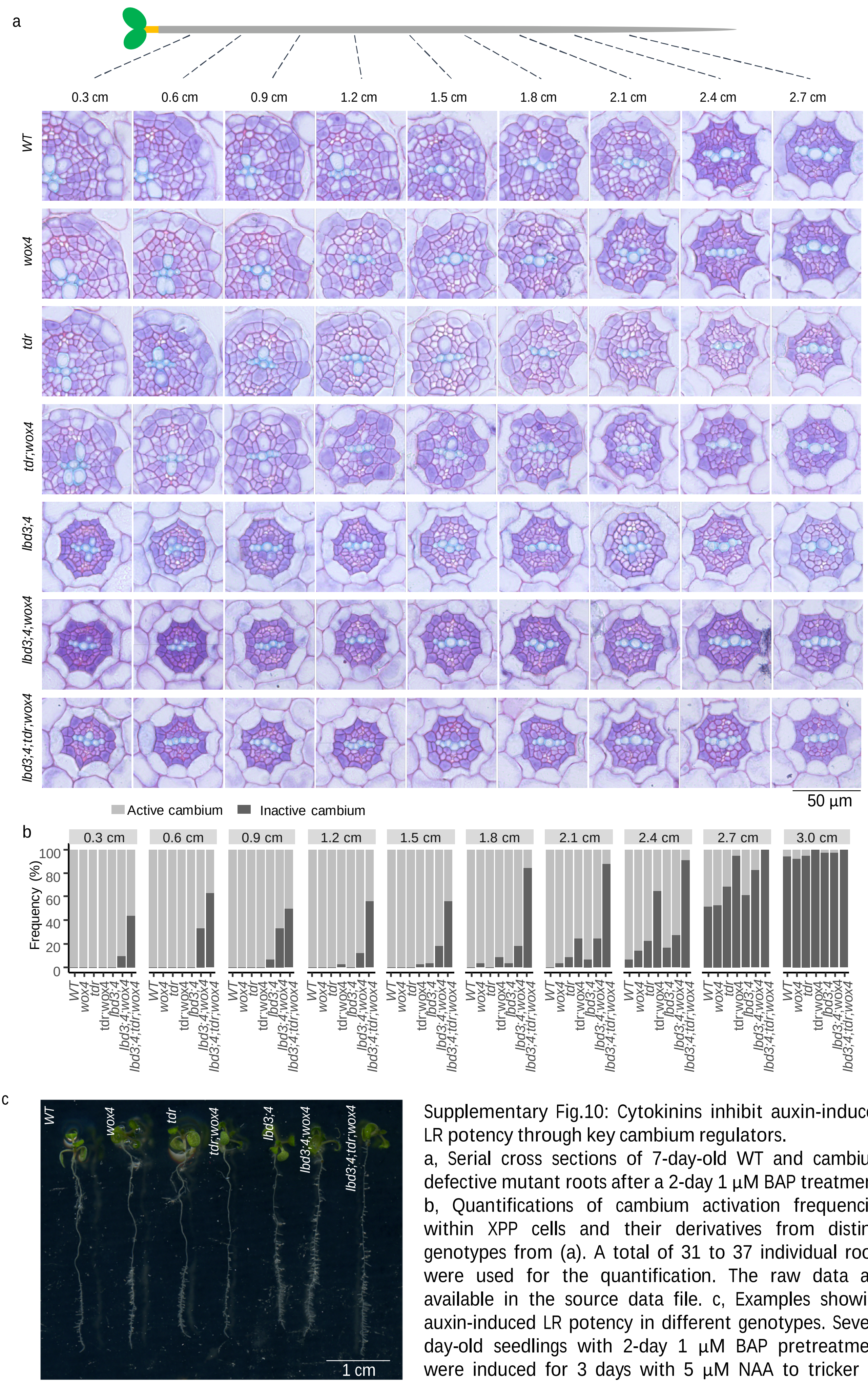


Supplementary Fig.10: Cytokinins inhibit auxin-induced LR potency through key cambium regulators.
a, Serial cross sections of 7-day-old WT and cambium defective mutant roots after a 2-day 1 µM BAP treatment. b, Quantifications of cambium activation frequencies within XPP cells and their derivatives from distinct genotypes from (a). A total of 31 to 37 individual roots were used for the quantification. The raw data are available in the source data file. c, Examples showing auxin-induced LR potency in different genotypes. Seven-day-old seedlings with 2-day 1 µM BAP pretreatment were induced for 3 days with 5 µM NAA to tricker LR formation.

Supplementary Table 1: Primers used in this study.

| Primer name | Sequence | Purpose |
|---|---|---|
| attB4-pHB53-F2 | ggggacaactttgtatagaaaagttgccgttcgttgcccacacattact | 1R4z-pHB53 |
| attB1-pHB53-R2 | ggggactgctttttgtacaaacttgctttctctctctagtttttcagac | |
| AscI-pHB53-F | ttcagaggcgcgccgttcgttgcccacacattact | pHB53-XVE |
| XhoI-pHB53-R | agcgtgctcgagtttctctctctagtttttcagac | |
| AscI-GATA23-F | ttcagaggcgcgccgaactcttgcctacctattaaaatcagc | pGATA23-XVE |
| FseI-GATA23-R | agcgtgggccggcccaaataaaaaaaaaacaatcttagttcttag | |
| AscI-35S-F | tatagaaaagttggcgcgccacgataccggtggtaccaacatggtg | 1R4a-AscI-35S-FseI-XVE |
| FseI-35S-R | aacgctttcattccgcccggaattggggccggcctacgtactcgaggtcctctcc | |
| XhoI-XPP-F | agactcgaggtgtttggttcgtaattatataccc | pXPP-XVE |
| XhoI-XPP-R | agactcgagtttggaaatcttcgtgtgttaagac | |
| erGUSplus-YFP-F | atttcggctacaagaacgctagccatcaccatcaccatcacgtggaattcggtggtggtggcggcggcgtgagcaagggcgaggagctg | 221z-erGUSpYFP |
| erGUSplus-YFP-R | caacaaattgataagcaatgctttcttataatgccaactttgtacaagaaagctgggtgttaaagctcatcatgcttgtacagctcgtccatgc | |
| 35Slong-loxp-F | gacctgttcgttgcaacaaattgataagcaatgctttttataatgccaactttgtatagaaaagttgatttaggtgacactatagaata | 1R4z-35Slong-Loxp |
| 35Slong-loxp-R | gaagttgcaaagctctctagctagaataacttcgtataatgtatgctatacgaagttataagcttgggcttgatatcactagtgcggccg | |
| HTR5-loxp-F | gacctgttcgttgcaacaaattgataagcaatgctttttataatgccaactttgtatagaaaagttgagatccgatataacaaaatttg | 1R4z-pHTR5-Loxp |
| HTR5-loxp-R | gaagttgcaaagctctctagctagaataacttcgtataatgtatgctatacgaagttataagcttgggctctgttcaacaaattaacaag | |
| MCS-35S-F | gacctgttcgttgcaacaaattgataagcaatgctttttataatgccaactttgtatagaaaagttgcctaggcatgcaattgactagtatcgtacccctactccaaaaatg | 1R4z-MCS-35S-MCS-Loxp |
| MCS-35S-R | gaagttgcaaagctctctagctagaataacttcgtataatgtatgctatacgaagttataagcttctcgaggtaccttcgaagtcctctccaaatgaaatg | |
| AvrII-UBQ10-F2 | gtacgctgggtcctagggtcgaaagcagcgatcaagc | 1R4z-pUBQ10-Loxp |
| Xho I-UBQ10-R | gtacgctgggtctcgagctgttaatcagaaaaactcagattaatc | |
| BSAI-gPXY-F | ttcagaggtctctctcgatgaaaaagaagaacatttctcc | 221z-PXY |
| BSAI-gPXY-R | agcgtgggtctcgggtgtcacaccccaatcctttgacatt | |
| YFP-F | ggggacaagtttgtacaaaaaagcaggctcgatggtgagcaagggcgagga | 221z-YFP-WOX4 |
| WOX4-YFP-R | gaaaactcatgaaccttcatccccccgccacccttgtacagctcgtccat | |
| YFP-WOX4-F | atggacgagctgtacaagggtggcggggggatgaaggttcatgagttttc | |
| WOX4-R | agcaaaccctaaacgcaactccaagtctaagtcaagtcatctcccttcaggatgga | |
| BsaI-pPER15-F | ttcagaggtctcttgcccaagctaaaatgtttttttatg | 1R4z-pPER15 |
| BsaI-pPER15-R | agcgtgggtctcgttgctgtcaaattttccgctagctag | |
| BSAI-LBD3-F | ttcagaggtctctctcgatgagacaaaagggtcacagacacg | 221z-LBD3 |
| BSAI-LBD3-R | agcgtgggtctcgggtggcaagaccaaaggaagtctccggc | |
| BSAI-LBD4-F | ttcagaggtctctctcgatgaaagaaagtagccggaagcaag | 221z-LBD4 |
| BSAI-LBD4-R | agcgtgggtctcgggtggcaagaccacatagactctccc | |
| BSAI-LBD11-F | ttcagaggtctctctcgatgctaaagatggagattaac | 221z-LBD11 |
| BSAI-LBD11-R | agcgtgggtctcgggtgtgtccaaagaggatcccaccac | |

Supplementary Table 2: Constructs used in this study.

| construct name | 1st box | 2nd box | 3rd box | destination vector |
|---|---|---|---|---|
| pPER15erYFP | 1R4z-pPER15 | 221z-erYFP | 2R3a-nosT | pHm43GW |
| pGATA23-XVE-CRE | 1R4a-pGATA23-XVE | 221z-CRE | 2R3a-nosT | pFRm43GW |
| pHB53-XVE-CRE | 1R4a-pHB3-XVE | 221a-CYCB1;1-CRE | 2R3a-nosT | pFRm43GW |
| pHB53-XVE-CYCB1;1-CRE | 1R4a-pHB3-XVE | 221a-CYCB1;1-CRE | 2R3a-nosT | pFRm43GW |
| pHB53-CYCB1;1-CRE | 1R4a-pHB3-XVE | 221a-CYCB1;1-CRE | 2R3a-nosT | pFRm43GW |
| pXPP-XVE-CYCB1;1-CRE | 1R4a-pXPP-XVE | 221a-CYCB1;1-CRE | 2R3a-nosT | pFRm43GW |
| 35S-loxp-erGUSpYFP | 1R4z-35S-loxp | 221z-erGUSpYFP | 2R3a-nosT | pBm43GW |
| 35S-loxp-erGUSpRFP | 1R4z-35S-loxp | 221z-erGUSpRFP | 2R3a-nosT | pBm43GW |
| pHTR5-loxp-erGUSpYFP | 1R4z-pHTR5-loxp | 221z-erGUSpYFP | 2R3a-nosT | pBm43GW |
| pHTR5-loxp-erGUSpRFP | 1R4z-pHTR5-loxp | 221z-erGUSpRFP | 2R3a-nosT | pBm43GW |
| pUBQ10-loxp-erGUSpYFP | 1R4z-pUBQ10-loxp | 221z-erGUSpYFP | 2R3a-nosT | pBm43GW |
| pUBQ10-loxp-erGUSpRFP | 1R4z-pUBQ10-loxp | 221z-erGUSpRFP | 2R3a-nosT | pBm43GW |
| pGATA23-XVE-LBD3-YFP | 1R4a-pGATA23-XVE | 221z-LBD3 | 2R3a-4xgly-YFP | pBm43GW |
| pGATA23-XVE-LBD4-YFP | 1R4a-pGATA23-XVE | 221z-LBD4 | 2R3a-4xgly-YFP | pBm43GW |
| pGATA23-XVE-LBD11-YFP | 1R4a-pGATA23-XVE | 221z-LBD11 | 2R3a-4xgly-YFP | pFRm43GW |
| pGATA23-XVE-ANT-YFP | 1R4a-pGATA23-XVE | 221z-ANT | 2R3a-4xgly-YFP | pFRm43GW |
| pGATA23-XVE-ATHB8-YFP | 1R4a-pGATA23-XVE | 221z-ATHB8 | 2R3a-4xgly-YFP | pFRm43GW |
| pGATA23-XVE-PXY | 1R4a-pGATA23-XVE | 221z-PXY | 2R3a-nosT | pFRm43GW |
| pGATA23-XVE-YFP-WOX4 | 1R4a-pGATA23-XVE | 221z-YFP-WOX4 | 2R3a-nosT | pFRm43GW |
| DR5erRFP | 1R4z-DR5 | 221z-erRFP | 2R3a-3AT | pHm43GW |
